# Human Neural Stem Cells Differentiate and Integrate, Innervating Implanted zQ175 Huntington’s Disease Mouse Striatum

**DOI:** 10.1101/2021.01.18.427078

**Authors:** Sandra M. Holley, Jack C. Reidling, Carlos Cepeda, Alice Lau, Cindy Moore, Iliana Orellana, Brian Fury, Lexi Kopan, Sylvia Yeung, Michael Neel, Dane Coleal-Bergum, Edwin S. Monuki, Charles K. Meshul, Gerhard Bauer, Michael S. Levine, Leslie M. Thompson

## Abstract

Huntington’s disease (HD), a genetic neurodegenerative disorder, primarily impacts the striatum and cortex with progressive loss of medium-sized spiny neurons (MSNs) and pyramidal neurons, disrupting cortico-striatal circuitry. A promising regenerative therapeutic strategy of transplanting human neural stem cells (hNSCs) is challenged by the need for long-term functional integration. We previously described that hNSCs transplanted into the striatum of HD mouse models differentiated into electrophysiologically active immature neurons, improving behavior and biochemical deficits. Here we show that 8-month implantation of hNSCs into the striatum of zQ175 HD mice ameliorates behavioral deficits, increases brain-derived neurotrophic factor (BDNF) and reduces mutant Huntingtin (mHTT) accumulation. Patch clamp recordings, immunohistochemistry and electron microscopy demonstrates that hNSCs differentiate into diverse neuronal populations, including MSN- and interneuron-like cells. Remarkably, hNSCs receive synaptic inputs, innervate host neurons, and improve membrane and synaptic properties. Overall, the findings support hNSC transplantation for further evaluation and clinical development for HD.

## Introduction

Huntington’s disease (HD) is a devastating neurodegenerative disorder that typically strikes individuals in midlife and progresses over 15-20 years before patients succumb to the disease (Ghosh and Tabrizi, 2018). HD is caused by an autosomal dominant CAG (glutamine) repeat expansion in the *huntingtin* (*HTT*) gene (The Huntington’s Disease Collaborative Research Group, 1993). Symptoms include progressive movement abnormalities, most notably chorea, difficulties with daily tasks, cognitive decline and psychiatric manifestations including depression, memory loss, and eventually dementia (Bates et al., 2002; Harper and Jones, 2002). Neuropathologically, the disease substantially impacts the striatum and cerebral cortex, with progressive loss of medium-sized spiny neurons (MSNs) and cortical pyramidal neurons, as well as loss of cortico-striatal synapses, leading to severe atrophy (Vonsattel and DiFiglia, 1998; Waldvogel et al., 2015). At the molecular level, the disease is accompanied by progressive loss of neuronal proteins, including brain-derived neurotrophic factor (BDNF) that supports survival of striatal neurons, as well as aberrant accumulation of aggregated huntingtin (HTT) protein species that correspond to disease pathogenesis (Saudou and Humbert, 2016). There is currently no FDA approved disease modifying treatments for HD patients that can either delay onset or modify disease progression. Recent strategies that show promise include DNA-targeting techniques such as zinc-finger proteins and CRISPR/Cas9, as well as HTT-lowering techniques currently in clinical trials, such as RNAi and antisense oligonucleotides (Tabrizi et al., 2019). However, these strategies also have limitations including efficient and targeted delivery, as well as the inability to replace or compensate for neuronal loss. Thus, there is an urgent need to finding additional therapeutic approaches.

In recent years, there has been an explosion of studies in regenerative medicine. The use of neural stem cells (NSCs) for the treatment of neurological disorders is in the early stages but there is already a wealth of information indicating that NSCs may offer a viable therapeutic avenue (El-Akabawy et al., 2012; Choi and Hong, 2017; Connor, 2018). We recently demonstrated that human (h)NSCs implanted in the striatum of R6/2 mice, a severe and rapidly progressing model of HD akin to juvenile HD (Mangiarini et al., 1996), survive, are functional, and improve a number of HD phenotypes (Holley et al., 2018; Reidling et al., 2018). Studies also included the long-lived full-length homozygous Q140 HD mouse model and we showed behavioral improvements and reduced aggregation; however, characterization of cells was limited due to low cell survival rate, perhaps caused by insufficient immunosuppression methods. Therefore, in the present study, we determined whether hNSCs can survive for longer periods of time, the types of cells they differentiate into in the host brain, whether cells are electrophysiologically active, whether they make connections with host cells and if neuroprotective effects persist. For these purposes, we used the heterozygous zQ175 mouse model that recapitulates aspects of adult-onset HD (Menalled et al., 2012). Heterozygous mice do not show overt behavioral symptoms until approximately 6 months of age and become fully symptomatic at 8-12 months (Heikkinen et al., 2012). Electrophysiological studies have demonstrated altered passive and active membrane properties of MSNs in symptomatic animals, as well as changes in synaptic activity (Heikkinen et al., 2012; Plotkin et al., 2014; Indersmitten et al., 2015; Southwell et al., 2016; Sepers et al., 2018). These functional alterations are associated with significant loss of neuron spines. Here we tested the viability, morphological and electrophysiological properties of hNSCs, as well as their potential therapeutic benefits in zQ175 mice. hNSCs were implanted in the striatum of pre-symptomatic mice (2.5 months), behavioral tests were performed for 8 months, electrophysiological tests began when the mice became fully symptomatic (10.5 months of age) and tissue was collected for immunohistochemical, biochemical, and morphological analyses. Our data show that implanted hNSCs survive, and a subset differentiate into mature MSNs and interneurons, establish connections with the host neurons, and rescue specific electrophysiological and behavioral phenotypes.

## Results

### ESI-017 hNSCs Transplanted Long-Term in zQ175 HD Model Mice Engraft and Differentiate

Our previous studies showed beneficial effects of hNSC implantation in R6/2 and Q140 model mice (Holley et al., 2018; Reidling et al., 2018). Here we wished to comprehensively evaluate whether hNSCs could survive for extended periods of time, whether they could further differentiate, and whether they could functionally compensate for loss of connectivity and neuronal function, which has not yet been described in a genetic model of HD. GMP-grade ESI-017 hNSCs (Holley et al., 2018; Reidling et al., 2018) were acquired as frozen aliquots (UC Davis), thawed, and cultured without passaging using the same media reagents as in the GMP facility. Mice were dosed by intrastriatal stereotactic delivery of 100,000 hNSCs per hemisphere at 2.5 months of age. To examine long-term survival of hNSCs, zQ175 mice were sacrificed at 10.5 months of age (8 months post-implant). The fate of the implanted cells was determined using IHC with markers for human cells, neural progenitor cells, post-mitotic neurons, astrocytes and oligodendrocytes. Implanted ESI-017 hNSCs survived and remained in the striatum with little migration away from the needle tract in most mice. When proliferative hNSCs are implanted into mice they are non-proliferative 8 months post-implant as indicated by a lack of staining for the proliferation marker Ki67 or when analyzed for the incorporation of the nucleotide analog EdU (**Fig. 1A & B**). Implanted hNSCs also did not express the neural stem cell marker nestin (**Fig. 1C**), supporting that they have differentiated. A survey of cell markers revealed that the implanted hNSCs differentiated into previously observed lineages of immature neurons (doublecortin, DCX+, **Fig. 1D & E**), very few astrocytes (glial fibrillary acidic protein, GFAP+, **Fig. 1C**), but not oligodendrocytes. Some cells appeared to differentiate into mature neurons (neuronal nuclei, NeuN+, **Fig. 1D, E & I** and BetaIII tubulin+, **Fig. 1G**) or interneurons (calretinin, CR+, **Fig. 1F**, or glutamic acid decarboxylase 65/67, Gad65/67+, **Fig. 1G**). We also detected glutamate transporter, vGlut1+ puncta surrounding implants potentially from cortical terminals (**Fig. 1F)**. hNSCs also differentiated into MSNs (dopamine- and cAMP-regulated neuronal phosphoprotein, DARPP32+ and B-cell lymphoma/leukemia 11B, Ctip2+ **Fig. 1H, I & J**). In addition, there is evidence that some hNSCs differentiated into cell types that exhibit inhibitory neuronal signals (gamma-aminobutyric acid, GABA+ **Fig. 1J**). These longer duration survival studies suggest that, given enough time, the hNSCs are no longer proliferative and can differentiate into post-mitotic neurons typically found in striatum.

**Figure 1:**
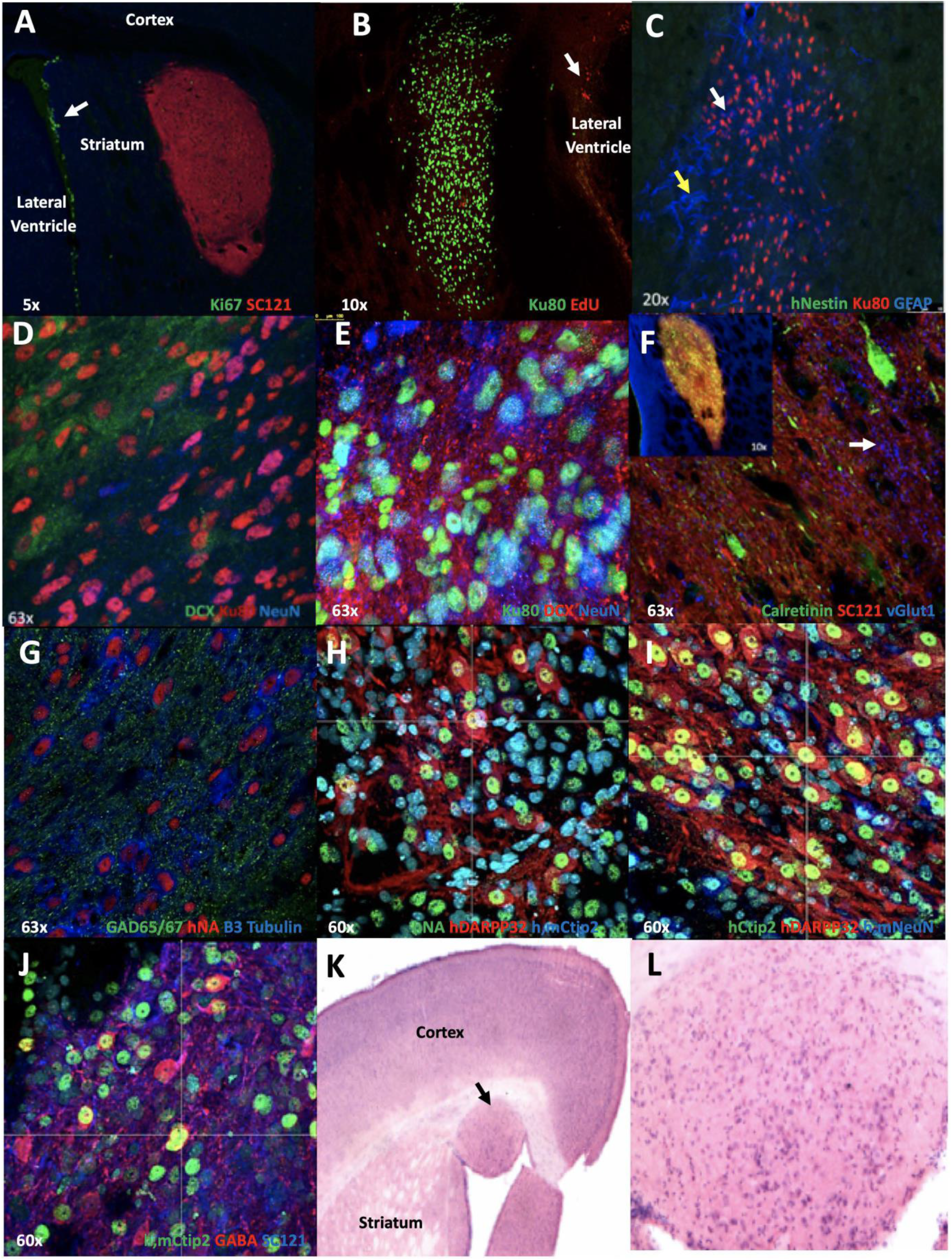
ESI-017 hNSCs implanted in zQ175 mice differentiate and do not proliferate: **A)** 5x mag. hNSCs (human cytosolic marker SC121, red) in zQ175 mice do not express the proliferation marker Ki67 (green). Proliferating cells in the lateral ventricle are indicated by arrow. **B)** 10X mag. showing hNSCs (human nuclear marker Ku80, green) did not incorporate the nucleotide analog EdU 24 hrs post injection indicating they are not dividing. **C)** 20x shows hNSCs (Ku80, red) do not express the neuronal progenitor marker nestin (green) but some cells show expression of the astrocyte marker GFAP (blue, white arrow). The hNSC implant site is surrounded by a mouse glial cell scar (GFAP+ blue, yellow arrow). **D)** hNSCs (Ku80, red) differentiate into both immature DCX+ (green) and more mature (NeuN, blue) neurons, shown at 63x. **E)** Another image of hNSCs (Ku80, green) differentiating into both immature DCX+ (red) and more mature (NeuN, blue) neurons, shown at 63x. **F)** 63x mag. hNSCs (SC121, red) differentiate into interneurons Calretinin (green) and some vGlut1 (blue, white arrow) puncta can be observed in the implantation site. The inset image shows the entire implant at 10x. **G)** 63x mag. hNSCs (Human nuclear antigen HNA, red) differentiate into a mixed population of cells that co-stain with GAD65/67 (green) or Beta III-tubulin (blue). **H)** Image shows hNSCs (HNA, green) differentiating into MSNs, DARPP32+ (red) Ctip2+ (blue) at 60x. **I)** Another 60x image showing hNSCs differentiating into MSNs using hDARPP32+ (red) and hCtip2+ (blue) only, as well as some other hNSCs expressing the mature neuronal marker NeuN. **J)** 60x image shows hNSCs (SC121, blue) expressing the MSN marker Ctip2 (green) and co-localization with the inhibitory neuronal marker GABA (red). **K)** 4x mag. and **L** (20x) showing H&E stains. A small nodule of cells that migrated away from the initial injection track are shown (at arrow, enlarged in L). Review from pathology included the comments that cytologically, the cells in the nodules are mostly well differentiated cells, with lots of large, mature-looking neurons.

After analysis of multiple brain sections, we observed evidence of some cell migration on the white matter tracts between the striatum and cortex (∼50% of the zQ175 but only a few WT mice). In addition, a subset of these mice that displayed cell migration (∼30% overall of implanted mice) exhibited nodules of cells that were positive for the human marker Ku80 adjacent to the cells in the implant site and in the ventricular space, but not in the striatum. H&E stains on adjacent sections of tissue (**Fig. 1K & L**) were performed and cytologically the cells in the nodules appeared to be mostly well-differentiated, mature-looking neurons. No evidence of proliferation was observed using Ki67 and nestin staining, suggesting these nodules are not a cause of concern in terms of potential for being or forming metastatic tumors. We performed IHC on another cohort of zQ175 and WT mice at 1, 2 and 5 months post implant in an attempt to observe the formation of nodules over time but did not observe any nodule formation. Between implantation of the original cohort and the nodule test mice we made a minor improvement to the surgical apparatus that may have altered hNSC migration on white matter tracks.

### ESI-017 hNSCs Improve Behavior in zQ175 HD Mice

We previously established that engrafted ESI-017 hNSCs significantly improve multiple behavioral outcomes in R6/2 and homozygous Q140 HD model mice (Reidling et al., 2018). To determine if hNSC implantation was also efficacious in heterozygous zQ175 mice used in this study we performed established behavioral assays for these mice. We found strikingly significant improvements in the running wheel test in mice that were 7.5 months old (5.5 months post implant) in ESI-017 hNSC implanted heterozygous zQ175 mice compared to vehicle mice, suggesting a reversion to WT levels and persistence of the effect **(Fig. 2A)**. The slope of motor learning was not significantly different among the 3 groups. In addition, we found significant improvements in distance traveled and velocity for hNSC-treated male and female mice combined compared to vehicle in the open field in mice that were 8 months of age (6 months post-implant) **(Fig. 2B & C)**. All open field behavioral outcomes are provided in Supplementary Materials **(Suppl. Fig. 1A-C)**.

**Figure 2:**
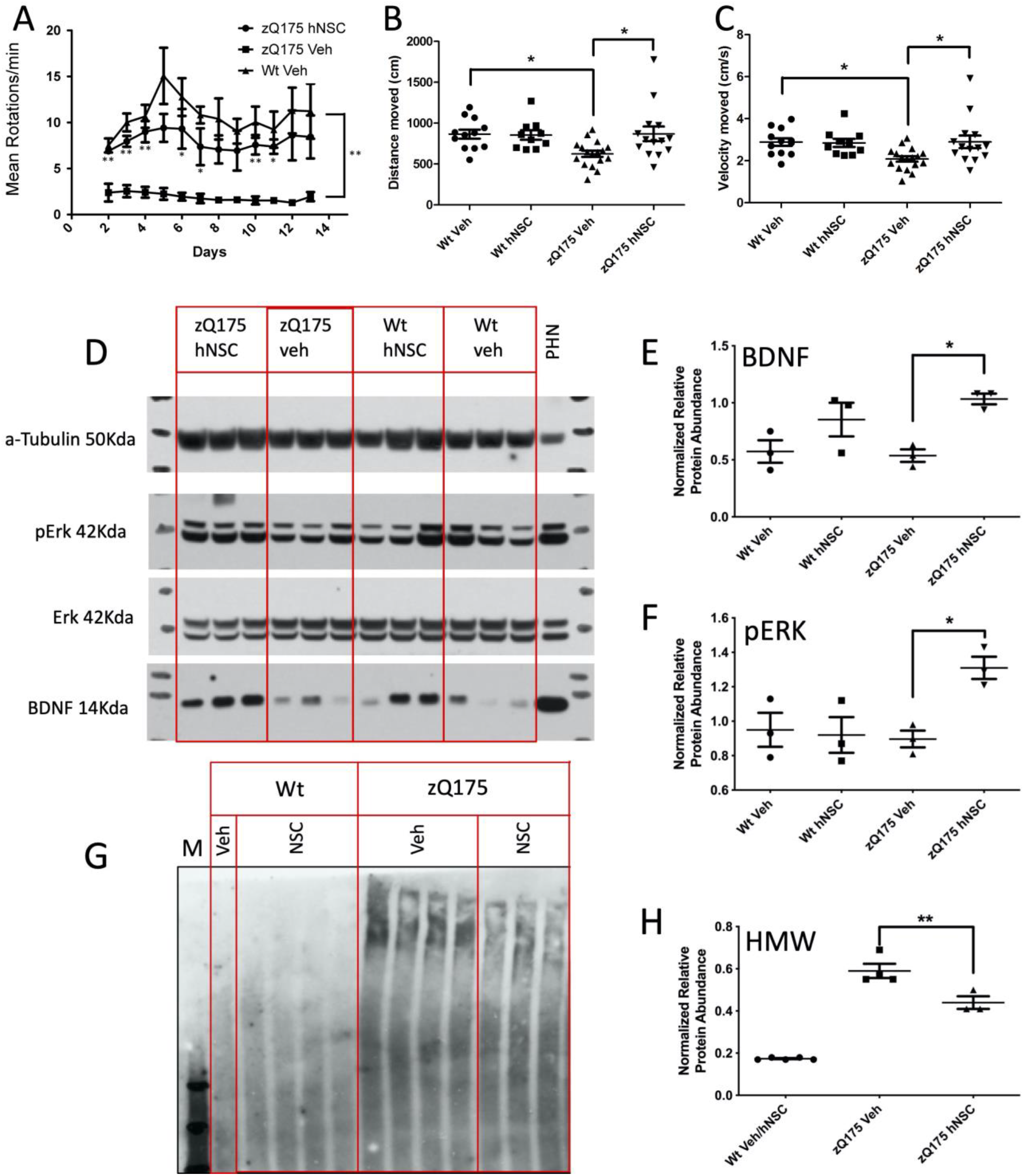
ESI-017 hNSCs implanted in zQ175 mice improve behavior, increase BDNF and p-ERK levels and reduce levels of a HMW mHTT species: **A**. Running wheel shows persistent improvement 5.5 months post-treatment. Mean running wheel rotations/minute/night over two weeks, in WT, zQ175 vehicle (veh) treated or zQ175 hNSC and zQ175 veh treated male mice (5/group). Results are expressed as the mean ± S.E.M one-way ANOVA Tukey HSD and Bonferroni post test p values for zQ175 veh treated to zQ175 hNSC treated Day2=0.003 Day3=0.001, Day4=0.002, Day5=0.06, Day6=0.03, Day7=0.03, Day8=0.07, Day9=0.06, Day10=0.009, Day11=0.01, Day12=0.08, Day13=0.1 summarized as * p<0.05, ** p<0.01, shown below error bars. WT veh to zQ175 veh Day2=0.001 Day3=0.001, Day4=0.001, Day5=0.002, Day6=0.002, Day7=0.001, Day8=0.005, Day9=0.01, Day10=0.001, Day11=0.002, Day12=0.02, Day13=0.03, summarized as **p≤0.03 at right. WT veh to zQ175 hNSC was not significantly different at any time-point. **B**. Total distance traveled in the open field 6 months post implant. Mice were subjected to the open field and total distance in centimeters of their respective tracks were combined and statistically analyzed to visualize any differences in ambulation. The zQ175 veh treated mice traveled less distance than WT veh and zQ175 hNSC treated mice. **C**. Velocity traveled in the open field 6 months post implant. Mice were subjected to the open field and velocity traveled in centimeters/sec of their respective tracks were combined and statistically analyzed to visualize any differences in time of ambulation. The zQ175 veh treated mice traveled slower than WT veh and zQ175 hNSC treated mice. Groups for open field included: 7 male zQ175 Het hNSC, 7 female Het hNSC, 9 male zQ175 Het veh, 8 female zQ175 Het veh, 5 male WT hNSC, 5 female WT hNSC, 6 male WT veh, 6 female WT veh. Results are expressed as the mean ± S.E.M with one-way ANOVA Tukey HSD and Bonferroni post test: * p=0.03 for genotype and p=0.02 for hNSC treatment in zQ175. **D**. Western blot analysis of whole tissue mouse striatal lysates from zQ175 and WT mice. zQ175 mice exhibited non-significantly reduced levels of BDNF compared to WT mice and levels were significantly increased with hNSC treatment. zQ175 mice also showed significantly increased levels of pERK compared to vehicle controls. Quantitation of the relative protein expression for BDNF (**E**) and pERK (**F**) is shown. **G**. Western blot analysis of mouse striatal lysates separated into detergent-soluble and detergent-insoluble fractions. zQ175 mouse striatum is enriched in insoluble accumulated mHTT compared to WT mice. hNSC implantation in zQ175 mice results in a significant reduction of insoluble HMW accumulated HTT. Quantitation of the relative protein expression for mHTT is shown in (**H**) and hNSC implantation results in a significant decrease of insoluble HMW accumulated mHTT. Graph values represent means ± SEM and Western blots were analyzed with ImageJ for quantification of BDNF (normalized to tubulin), ERK/pERK ratio or aggregate type/section. Data was analyzed by one-way ANOVA with Tukey HSD and Bonferroni post hoc test (n = 3/group) ∗p = 0.03, ∗∗p = 0.005.

### Engrafted ESI-017 hNSCs Correlate with Increased BDNF and Decreased Pathogenic Accumulation of mHTT Proteins

Increased levels of BDNF were demonstrated after hNSC implantation in the rapidly progressing R6/2 HD mouse model (Reidling et al., 2018), therefore we evaluated whether this effect could be sustained following long-term engraftment in zQ175 mice. Striatal BDNF quantified by Western blot analysis was slightly but not significantly decreased in a subset of male zQ175 mice (n=3/group) compared to WT, but a significant increase in BDNF levels was observed in hNSC-treated zQ175 mice compared to vehicle **(Fig. 2D & E)**. Interestingly, we also observed an increase in the phosphorylation of extracellular signal-regulated kinase (ERK) protein suggesting potential activation of cellular signaling cascades **(Fig. 2D & F)**. In addition, our previous studies showed that hNSC treatment can reduce high molecular weight (HMW) mHTT species, a pathogenic marker for HD. Consistent with those results we also observed persistent reduction in levels of a HMW mHTT species in hNSC treated zQ175 mice **(Fig. 2G & H)**, suggesting prevention of pathology by the transplanted cells.

### Electrophysiological and Morphological Characterization of hNSCs

hNSCs transplanted for 4 weeks in R6/2 mice showed properties of immature electrophysiologically active hNSCs (Reidling et al., 2018), however, we did not know whether this could be sustained or improved in long-term implanted mice. To perform electrophysiological studies we used a subset of female zQ175 and WT mice (10.5 month-old) implanted at UCI and shipped live to UCLA. The hNSC grafts in zQ175 mice were easily identifiable under IR-DIC microscopy **(Fig. 3A1 & A2)**. In contrast to host tissue, which appeared darker due to myelin from fiber tracks, the graft appeared more translucent and densely populated by diverse cell types. In agreement with histological findings, most cells (∼80%) within the graft sites were small (<15 µm in diameter), round or bipolar, and had few extended processes. These cells appeared to be visually similar to the immature neuronal cell types previously recorded in R6/2 mice 4-6 weeks after implantation (Holley et al., 2018; Reidling et al., 2018). We also observed a number of cells that were larger in size (15-25 µm in diameter) with abundant and extensive processes that were visually different from host MSNs. As immature-looking hNSCs were characterized extensively in our previous publication, in the present study we focused our recordings on these larger, more mature-looking cells. We found for all hNSC types, membrane properties were similar for cells recorded in either zQ175 or WT mice and data from recorded hNSCs from both genotypes were pooled **(Table IA, Fig. 3C)**.

**Figure 3:**
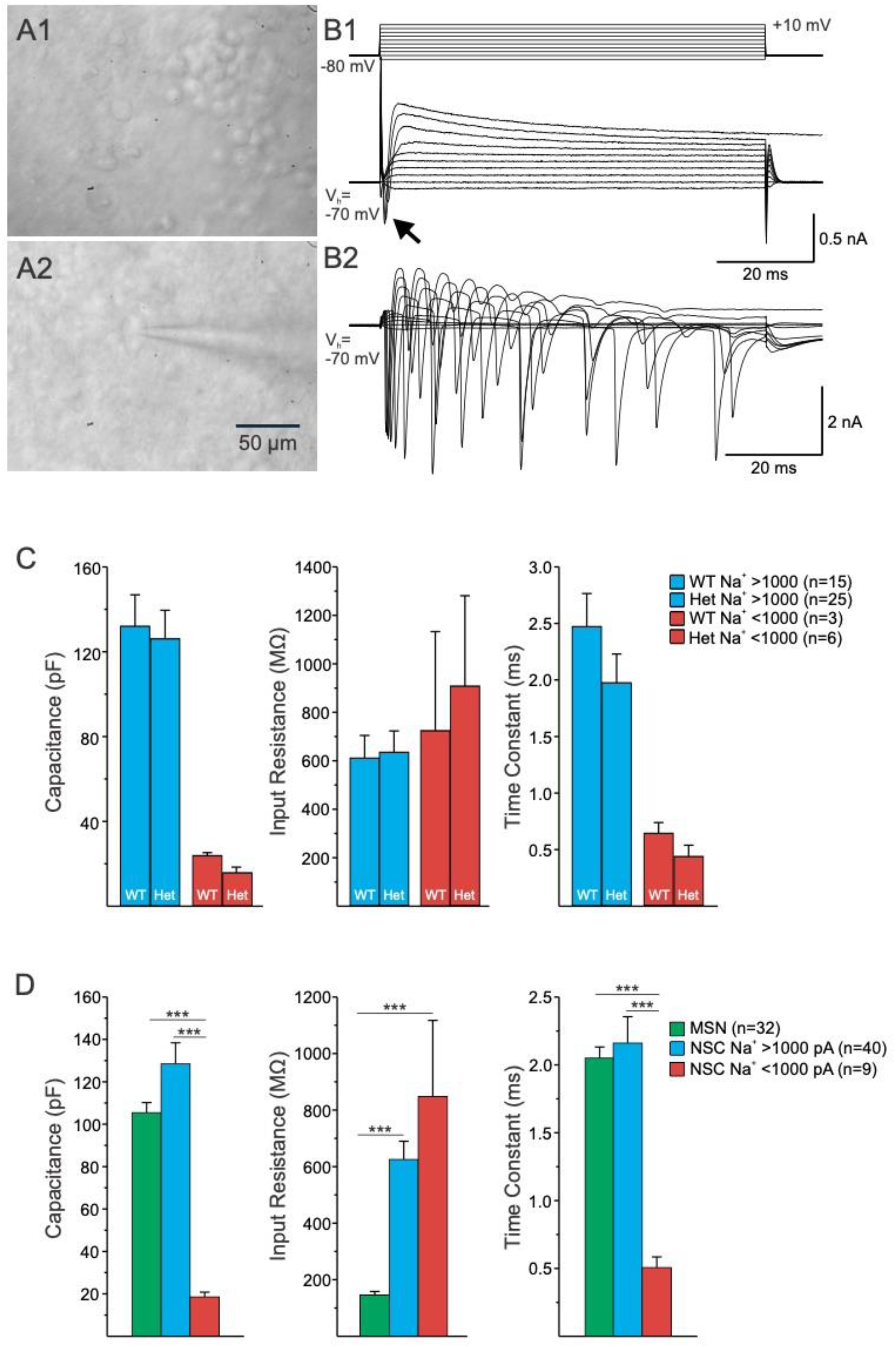
Two main types of hNSCs could be distinguished based on soma size, passive and active membrane properties. **(A1)**. IR-DIC image from transplanted hNSCs in a zQ175 mouse revealed that most cells had small, round somata. **(A2)**. A small percentage of hNSCs had larger somata and some dendritic branches. A patch pipette attached to the cell can be seen. Example cell is from a zQ175 mouse. **(B1)**. Recordings performed on hNSCs in zQ175 mice show that depolarizing step voltage commands from a holding voltage (V_h_) of −70 mV induced only incipient Na^+^ currents (<1000 pA) and large outward K^+^ currents. These cells were deemed immature. Example trace is from a zQ175 mouse. Voltage clamp protocol is shown above the current traces. **(B2)**. Large cells displayed repetitive large Na^+^ spikes (>1000 pA) and were deemed mature. Example trace is from a WT mouse receiving an implant of hNSCs. **C**. Bar graphs show mean±S.E.M of cell membrane properties of two types of implanted hNSCs, those with immature (red) and those with mature (blue) membrane properties, using Na^+^ current amplitude as cut-off. Capacitance, input resistance and time constants are shown for recorded hNSCs. **D**. Bar graphs show mean±S.E.M of cell membrane properties of host MSNs (WT and zQ175 pooled together) compared to implanted hNSCs. Capacitance, input resistance and time constants are shown for recorded host MSNs and mature and immature hNSCs. Statistical significance was measured using one-way ANOVA tests followed by Bonferroni *post hoc* tests for pairwise comparisons and *** represents p<0.001. In (D.) Capacitance: p=2.0007e-05 for MSN versus NSC Na+<1000 pA and p=6.1282e-08 for NSC Na+>1000 pA versus NSC Na+<1000 pA. Input Resistance: p= 5.3000e-06 for MSN versus NSC Na+>1000 pA and p= 2.6645e-05 for MSN versus NSC Na+<1000 pA. Time Constant: p=8.9566e-05 for MSN versus NSC Na+<1000 pA and p=1.7886e-05 for NSC Na+>1000 pA versus NSC Na+<1000 pA.

Recorded cells in the grafts appeared to differentiate into two groups, one population of cells with immature-like electrophysiological properties and one population with more mature-like properties. In total, 49 hNSCs were recorded (n=18 in WT and 31 in zQ175 mice). The two groups of hNSCs were separated based on whole-cell patch-clamp recordings in voltage clamp mode measuring passive membrane properties (cell membrane capacitance, input resistance and decay time constant), as well as Na^+^ current amplitudes. Cells with low membrane capacitance (mean±s.e., 18.6±2.7 pF), high input resistance (847.3±269.9 MΩ), fast time constant (0.51±0.08 ms), and Na^+^ current amplitude <1 nA were immature-looking hNSCs (n=9, 3 in WT and 6 in zQ175 mice) **(Fig. 3B1 & C)**. A number of these cells did not display Na^+^ currents and were probably glial cells, e.g., astrocytes (n=2 in WT and 5 in zQ175 mice). In contrast, cells with high membrane capacitance (128.5±10.0 pF), medium-high input resistance (625.1±64.6 MΩ), slower time constant (2.2±0.2 ms), and Na^+^ current amplitude >1 nA were mature-looking hNSCs (n=40, 15 in WT and 25 in zQ175 mice, **Fig. 3B2 & C)**.

To further characterize maturation and differentiation of hNSCs, we compared their basic membrane properties with those of MSNs recorded from the host **(Fig. 3D)**. While subtle differences in membrane properties have been reported in MSNs giving rise to the direct and indirect pathways (Cepeda et al., 2008; Gertler et al., 2008), in the present study we did not differentiate these two classifications of MSNs. Cells with Na^+^ currents >1 nA had large membrane capacitance, similar to or even larger than that of host MSNs. In contrast, cells with low amplitude Na^+^ currents had very low membrane capacitance, similar to recordings from immature neurons. Both types of hNSCs had relatively high membrane input resistance compared with MSNs. In addition, differences in decay time constants were similar to those observed for cell capacitance. About 50% of recorded hNSCs lacked inward Ca^2+^ currents, usually visible with Cs^+^-based internal solution after depolarizing voltage commands, suggesting that these cells were not projection MSNs. However, the remaining recorded hNSCs (n=10 in WT mice and n=14 in zQ175) displayed inward Ca^2+^ currents. Some of these cells displayed Ca^2+^ currents larger in amplitude and similar to those observed in host MSNs (n=5 in WT mice and n=6 in zQ175) **(Fig. 4)**.

**Figure 4:**
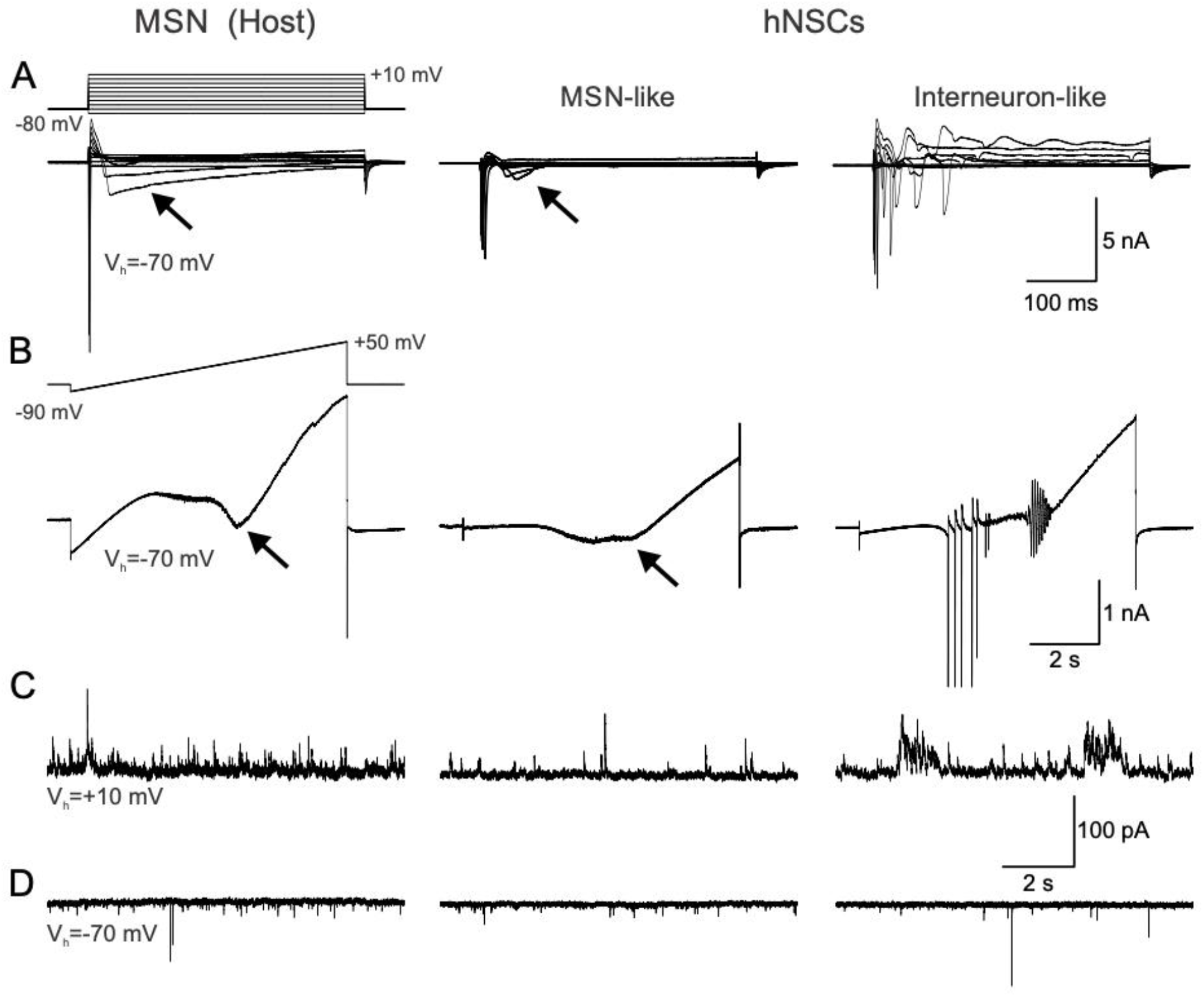
Examples of host MSN and mature hNSCs from a zQ175 mouse. Some hNSCs displayed mature neuronal properties that were similar to (MSN-like) and different (interneuron-like) from host MSNs. **A**. Recordings of intrinsic currents in response to step voltage commands (10 mV steps from −80 to +10 mV) in a host MSN (left panel) and in a MSN-like (center panel) and an interneuron-like (right panel) hNSC. Depolarizing voltage commands induced large Na^+^ currents followed by inactivating Ca^2+^ currents (arrows) of variable amplitudes. In the interneuron-like cell, repetitive spikes were observed but no prominent Ca^2+^ currents. **B**. Recordings of currents in response to a ramp voltage command (8 sec, from −90 to +50 mV). Some hNSCs recorded (center panel) had properties similar to host MSNs, both displaying Ca^2+^ currents (black arrows) after membrane depolarization. Other large hNSCs recorded lacked Ca^2+^ currents but displayed repetitive Na^+^ spikes (right panel) and were probably interneurons. **C**. Spontaneous synaptic currents recorded at +10 mV in a host MSN and in MSN-like and interneuron-like hNSCs. These currents are mostly GABAergic. **D**. Spontaneous synaptic currents recorded at −70 mV in a host MSN and in a MSN-like and an interneuron-like hNSC. These currents are most likely glutamatergic. Traces in each column are from the same cell. Calibrations on the right apply to all traces in each row.

The intrinsic membrane properties of these hNSCs were not significantly different regardless of the mouse genotype **(Table IB** & **C)**. These cells also displayed frequent spontaneous synaptic activity and could represent projection neurons with the potential to connect with other cells inside and outside the graft. Biocytin labeling revealed these cells had abundant dendritic processes and sparse spines **(Fig. 5)**.

**Table 1.**
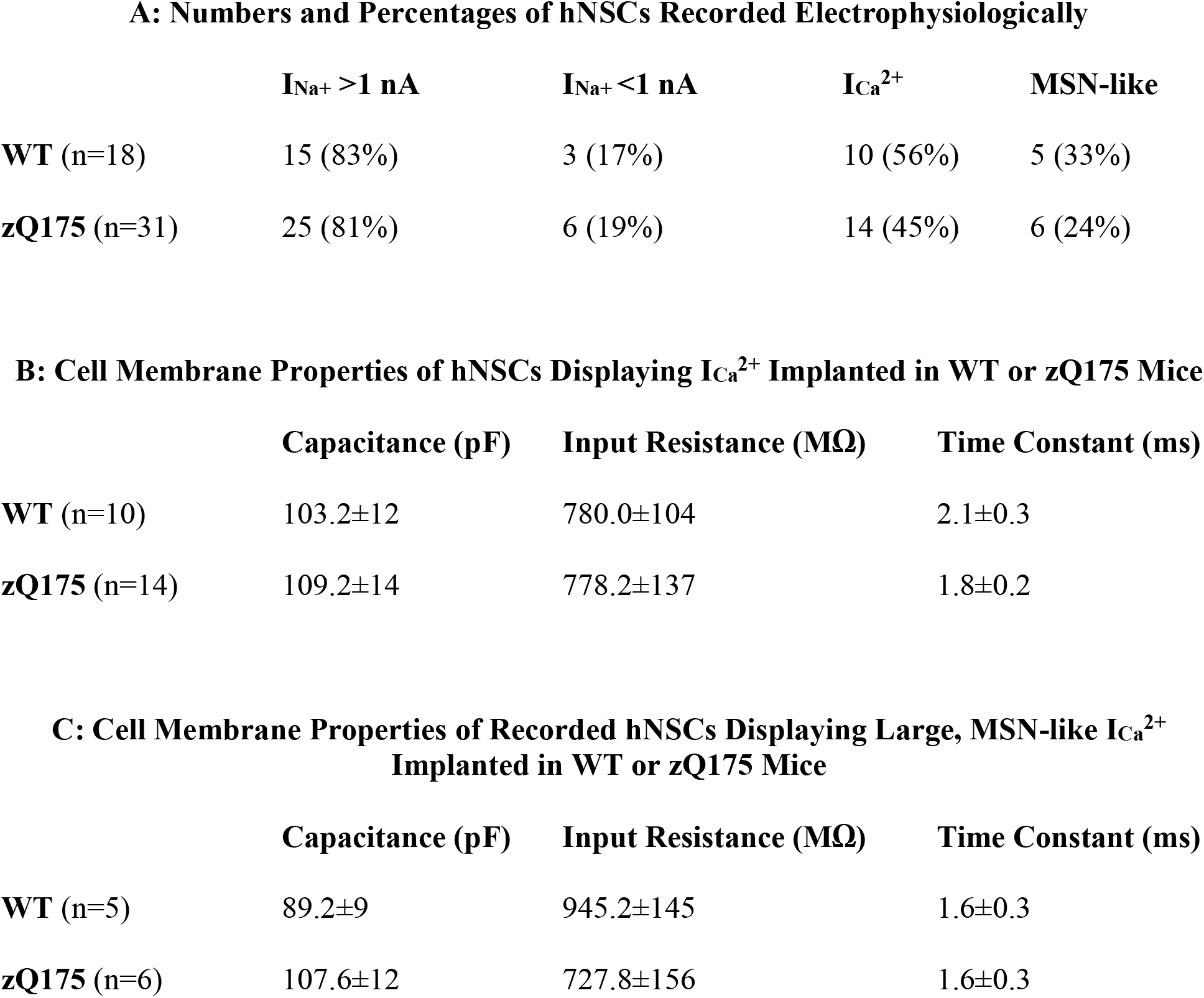
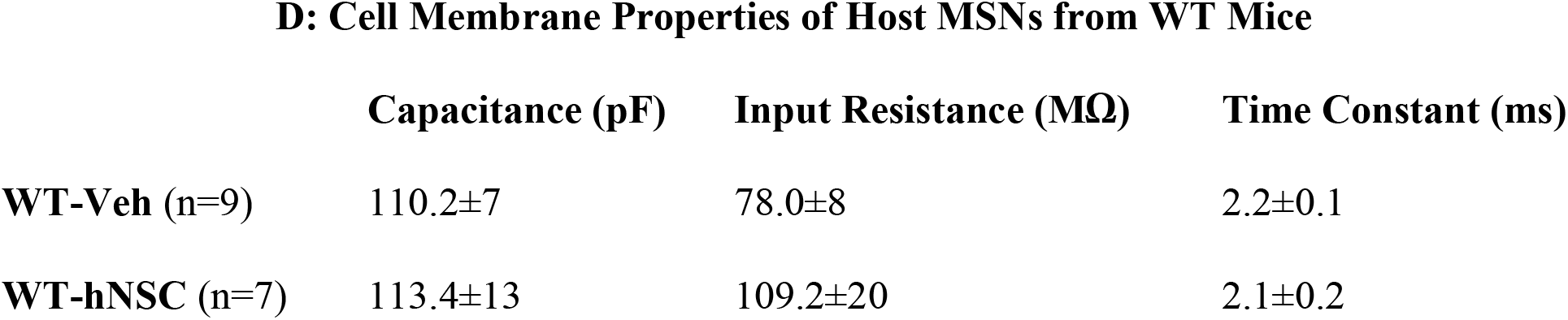

**Figure 5:**
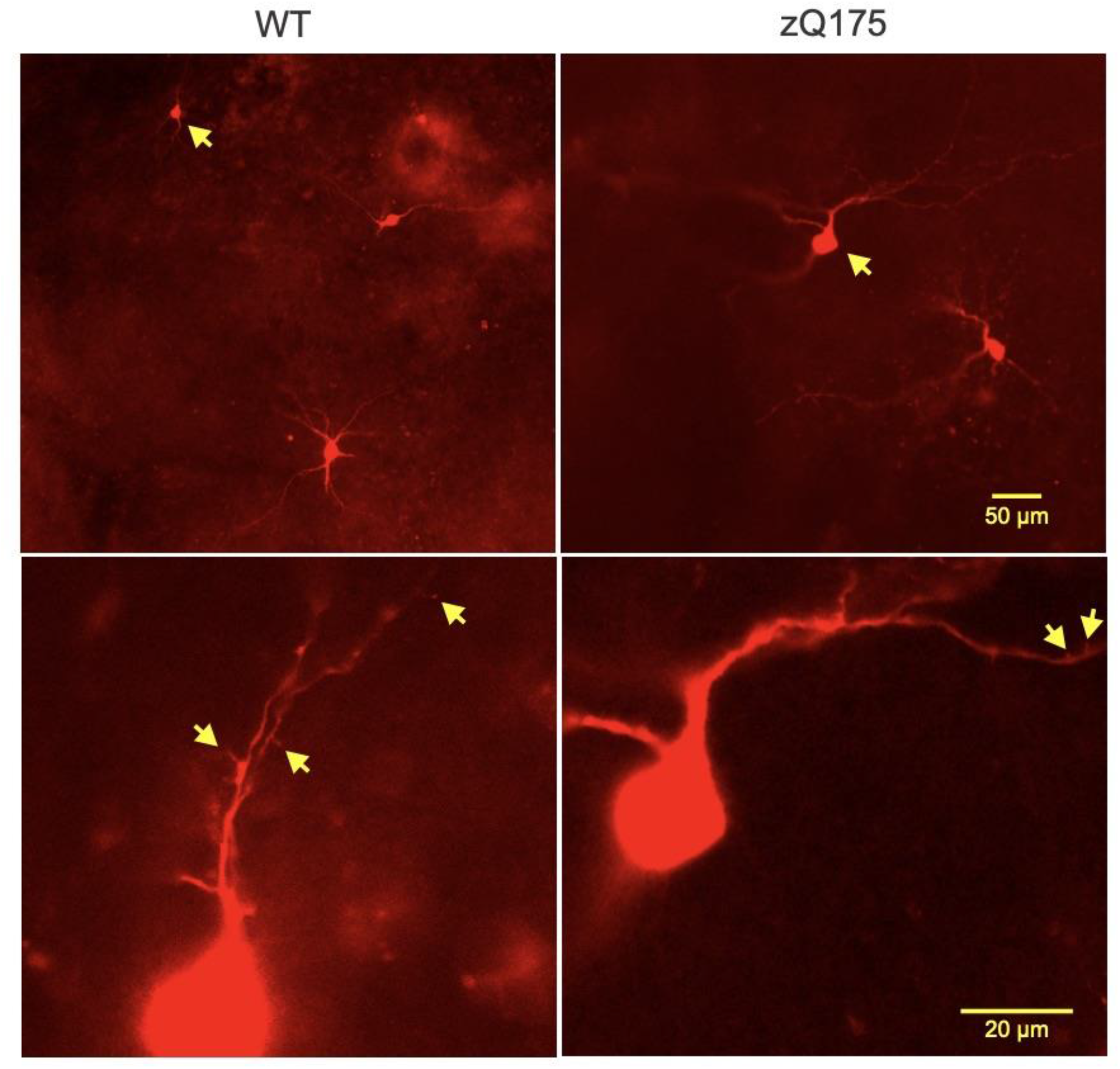
Biocytin-filled hNSCs recorded in WT and zQ175 mice in host striatum. Upper panels show low-magnification images of several hNSCs. Arrows indicate two cells that displayed MSN-like electrophysiological properties. Lower panels show processes from the same cells at higher magnification. Arrows indicate possible dendritic spines.

The remainder of hNSCs did not display Ca^2+^ currents. Instead, they displayed large inward Na^+^ and outward K^+^ currents, and could fire repetitive action potentials. We were able to recover ∼50% of biocytin-filled hNSCs (9/18 or 50% and 15/31 or 48% recorded in WT and zQ175 mice, respectively). Importantly, over 80% of those recovered clearly co-immunostained with the SC121 and Ku80 markers (8/9 or 89% and 12/15 or 80% in WT and zQ175 mice, respectively), thus confirming their human origin. Post-recording immunostaining of biocytin-filled large hNSCs with SC121 and Ku80 (with strepavidin-Alexa 594 for cell visualization) in fixed slices revealed hNSCs with relatively large somata (compared to host MSNs) and extensively branched processes. Other visible hNSCs were smaller in size and had either many or a small number of processes. The large hNSCs had cell diameters of up to ∼25 µm and were positive for both SC121 and Ku80 immunostaining (**Fig. 6**). Another important difference was that, compared with host MSNs or MSN-like hNSCs, many large hNSCs fired spontaneously. In cell-attached recording mode, cells fired rhythmically and also received rhythmic GABAergic synaptic inputs. We tentatively concluded these inputs were GABAergic because the GABA reversal potential was around −60 mV **(Suppl. Fig. 2)**. Due to this rhythmic input, one possibility is that the source of this GABA input is from other mature hNSCs, as hNSCs remained in close proximity. In addition, host GABAergic interneurons, such as the somatostatin-expressing or neuropeptide Y (NPY)-expressing interneurons that fire spontaneously, may contribute to these rhythmic inhibitory events recorded in hNSCs. Another type of recorded hNSC (n=3) resembled low-threshold spiking (LTS) striatal interneurons **(Suppl. Fig. 3A)**. In current clamp mode these cells displayed prominent delayed rectification at hyperpolarized membrane potentials **(Suppl. Fig. 3B)**. When depolarized they discharged in bursts of action potentials, seemingly riding on a low-threshold Ca^2+^ spike followed by a membrane hyperpolarization, which produced spontaneous oscillations and bursts of action potentials **(Suppl. Fig. 3C)**. The last type of hNSC (n=2) resembled cholinergic (ChAT-expressing) interneurons, also known as striatal tonically active neurons (TAN). They displayed rhythmic firing (2-3 Hz) and prominent delayed inward rectification. Electrophysiological identification of these interneurons was supported by IHC detection of appropriate markers **(Suppl. Fig. 4 and 5)**.

**Figure 6:**
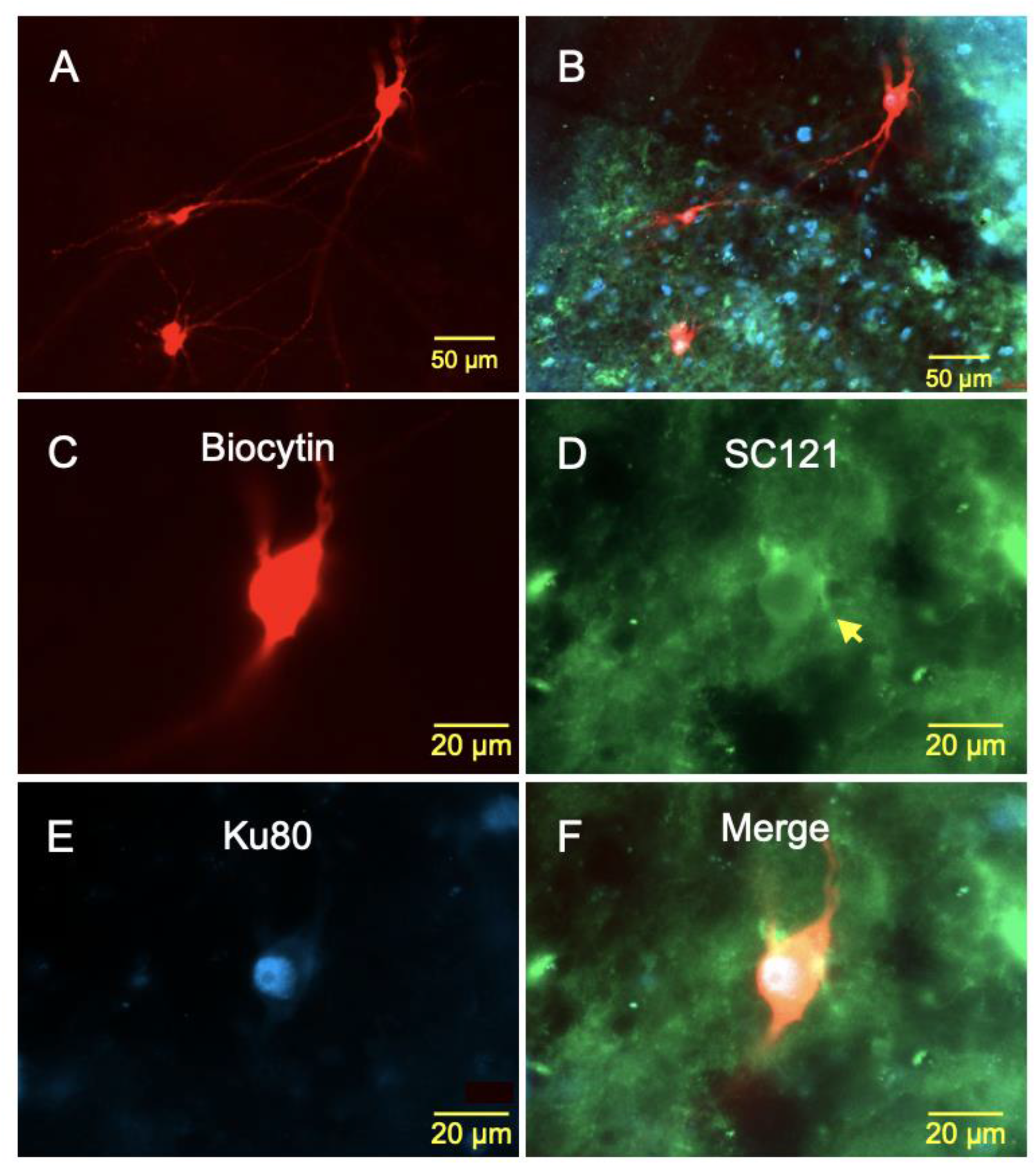
Images of large hNSCs from a zQ175 mouse, which displayed interneuron-like electrophysiological properties. **A**. Four medium to large hNSCs were recorded within the graft. **B**. The same image showing staining for the human markers SC121 (cytoplasmic, green) and Ku80 (nuclear, blue). **C**. Fluorescent image of one of the large biocytin-filled hNSCs. Immunostaining of the same hNSC with human stem cell markers SC121 (arrow in **D)** and Ku80 **(E). F**. Merged image showing biocytin and the two human stem cell markers. The electrophysiology of this interneuron-like hNSC is shown in Fig. 4 (right panels).

### Synaptic Properties of hNSCs Compared with Host MSNs

Glutamatergic inputs onto hNSCs were examined by holding the membrane at −70 mV. GABAergic inputs were examined by holding the membrane at +10 mV. Immature hNSCs displayed very few synaptic inputs whereas mature hNSCs had a wide range of synaptic inputs, some with frequencies as high as those recorded from MSNs [2.1±0.2 Hz (range 0.2-6.9 Hz) for MSNs versus 1.5±0.2 Hz (range 0.0-8.7 Hz) for hNSCs]. Based on the frequency of spontaneous synaptic activity, cells displaying Na^+^ currents could be divided into those with high IPSC and high EPSC frequencies, high IPSC and low EPSC frequencies, and low IPSC and high EPSC frequencies. As expected, large hNSCs displayed higher frequencies than those of immature-looking hNSCs **(Fig. 7A-C)**. Some hNSCs (n=2) also were tested for their ability to respond to electrical stimulation in the vicinity of the graft. Both cells responded to the electrical stimulation displaying both glutamatergic and GABAergic responses, as demonstrated by specific blockade with appropriate receptor antagonists **(Fig. 7D)**. Thus, these studies provided evidence that hNSCs establish synaptic contacts with the host and probably among hNSCs as well.

**Figure 7:**
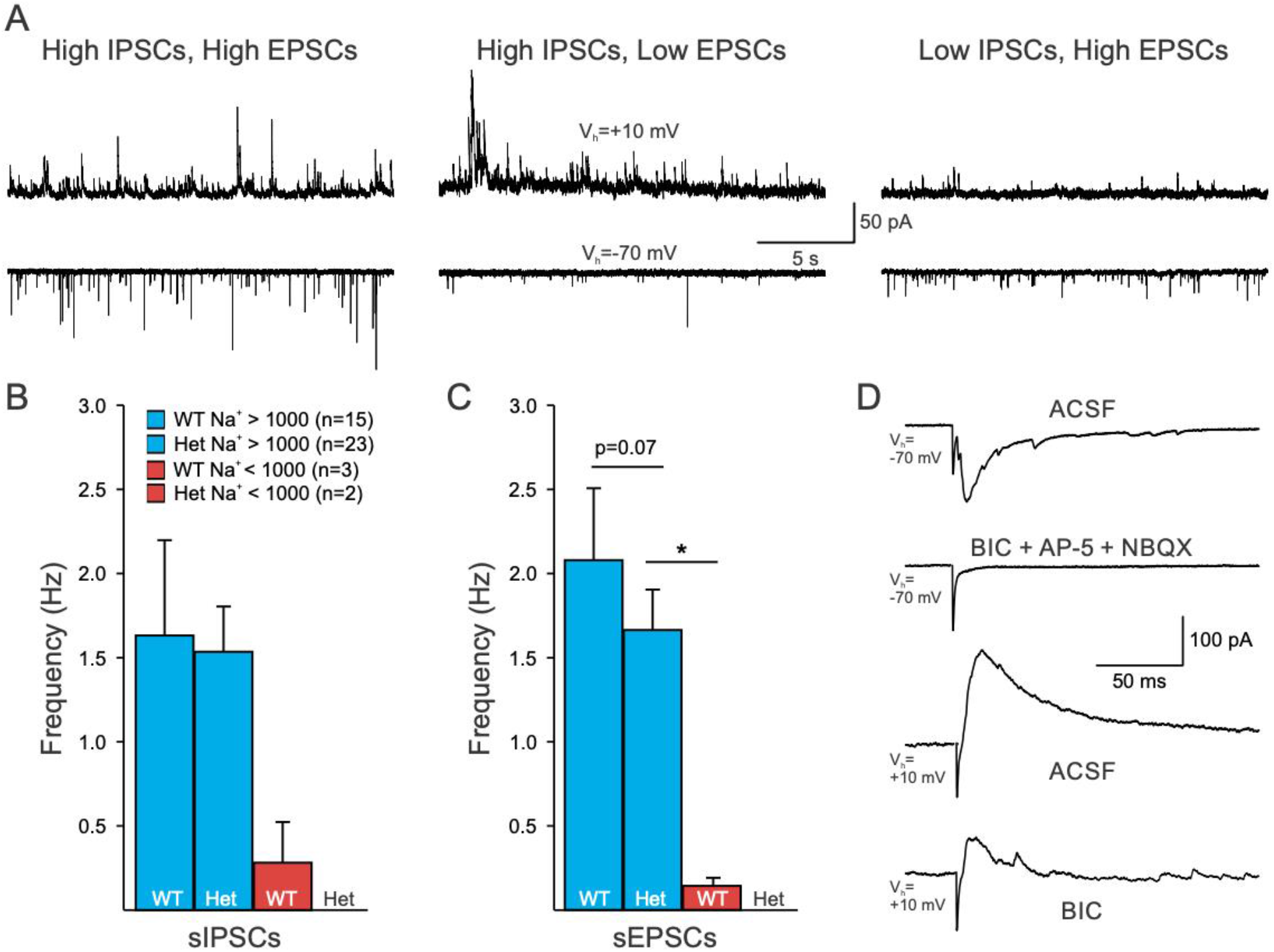
hNSCs in WT and Q175 mice display multiple synaptic inputs. **A**. Sample traces of sIPSCs and sEPSCs recorded from hNSCs in a zQ175 mouse in ACSF. **B** and **C**. Summary of the sIPSC and sEPSC frequencies from hNSCs categorized by the size of their Na^+^ currents. Statistical significance was measured between groups using Kruskal-Wallis one-way ANOVA on ranks followed by Holm-Sidak pairwise comparisons, where * p=0.024 for Het Na^+^>1000 pA versus WT Na^+^<1000 pA. **D**. Responses evoked by electrical stimulation (0.1-0.5 mA, 1 ms duration) of host striatal neurons (about 200 μm lateral to the graft) in a hNSC from a zQ175 mouse. Glutamatergic (V_h_=-70 mV) and GABAergic (V_h_=+10 mV) responses were reliably evoked by electrical stimulation. Responses were blocked by glutamate and GABA_A_ receptor antagonists respectively.

### Ultrastructural Evidence that hNSCs Establish Synaptic Contacts Within and Outside the Graft

To examine the morphology of implanted hNSCs and determine whether synaptic contacts are present between the host and transplanted hNSCs, thus further supporting the electrophysiology results, we performed electron microscopy (EM). EM studies provided additional anatomical evidence that hNSCs in the graft received innervation from host neurons. A subset of the mice implanted at UCI (3 females per group, zQ175 hNSC implanted) were sent live to the Portland VA Medical Center where the mice were perfused with fixative, the brains were collected and fixed for EM tissue processing. Results indicate that mouse host cell nerve termini make both symmetrical and asymmetrical synaptic contacts with implanted ESI-017 hNSCs. As a proof-of-principle, in a subset of samples we performed double-immunostaining for vGlut1 (which labels cortical glutamate terminals) and SC121. There were mouse host cell nerve termini that were positively labeled with vGlut1 making asymmetrical synaptic contacts with ESI-017 hNSCs **(Fig. 8A)**, suggesting that mouse (host) cortical neurons contribute to these connections. However, it is also possible that the other major host nerve terminal input to the ESI-017 hNSC implanted neurons, where an asymmetrical synaptic contact is observed **(Fig. 8B, C)**, may be from the thalamus.

**Figure 8:**
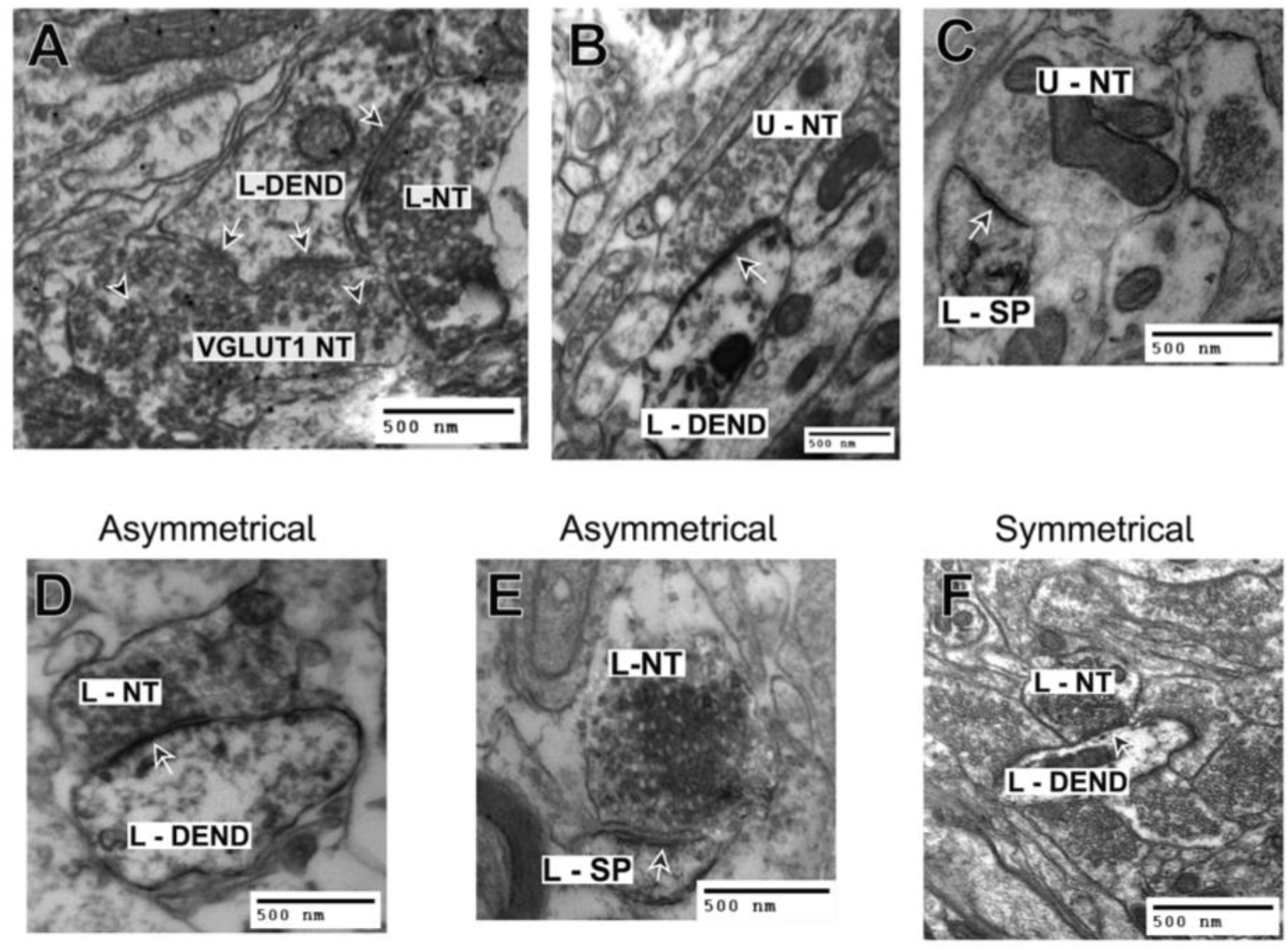
EM studies revealed that the host tissue makes synaptic contacts with ESI-017 hNSCs in zQ175 mice within the implantation site: **A**. hNSC/DAB labeled nerve terminal (L-NT) making a symmetrical synaptic contact (single black arrow with white outline) with an hNSC/DAB labeled dendrite (L-DEND). Vglut1 labeled nerve terminal (VGlut1 NT) making an asymmetrical synaptic contact with the head of a stubby spine that contains a perforated postsynaptic density (two black arrows with white outline pointing to synaptic contact). The VGlut1 NT (black arrowheads with white outlining-VIP labeling), is distinguished from the adjacent nerve terminal (which is DAB labeled). The primary origin of VGLUT-1 containing neurons is the cortex. **B**. An unlabeled nerve terminal (U-NT) making an asymmetrical synaptic contact (arrow) with an underlying hNSC positive labeled dendrite (L-DEND) (note the darkened DAB reaction product within the dendrite). The unlabeled nerve terminal might originate from either the host striatum, thalamus or cortex (see panel A), while the labeled dendrite originates from the implanted stem cells. **C**. An unlabeled nerve terminal (U-NT) is making an asymmetrical synaptic contact (arrow) with an underlying hNSC positive/labeled dendritic spine (L-SP) (note the darkened DAB reaction product within the spine). As in panel B, the unlabeled nerve terminal might originate from either the host striatum, thalamus or cortex (see panel A), while the labeled dendritic spine originates from the implanted stem cells. **D**. A labeled nerve terminal (L-NT) is making an asymmetrical synaptic contact (arrow) with an underlying hNSC positive/labeled dendrite (L-DEN) (note the darkened DAB reaction product within the nerve terminal and dendrite). The labeled nerve terminal and dendrite originate from the implanted stem cells. **E**. A labeled nerve terminal (L-NT) is making an asymmetrical synaptic contact (arrow) with an underlying hNSC positive/labeled dendritic spine (L-SP) (note the darkened DAB reaction product within the nerve terminal and spine). The labeled nerve terminal and spine originate from the implanted stem cells. **F**. A labeled nerve terminal (L-NT) is making a symmetrical synaptic contact (arrow) with an underlying hNSC positive/labeled dendrite (L-DEN) (note the darkened DAB reaction product within the nerve terminal and dendrite). The labeled nerve terminal and dendrite originate from the implanted stem cells.

Within the implant, of the host (non-labeled) terminals contacting SC121+ dendrites/spines in the implant site, 60% of the asymmetrical contacts were on dendrites and 40% on spines (total of 35 observations), suggesting contacts with MSN-like hNSCs. Of the host contacting SC121+ labeled dendrites, if the synaptic contact was symmetrical, 100% of those contacts were on the dendrite (22 observations). Of all the host (non-labeled) contacts onto SC121+ labeled cells in the implant area, 62.1% were asymmetrical while 37.9% of the contacts were symmetrical (total of 58 observations).

Within the implant, the percentage of SC121+ nerve terminals contacting SC121+ dendrites was determined. We found that of those total synaptic contacts, 71.3% were making an asymmetrical synaptic contact, with 27.8% making a symmetrical contact (total of 115 observations). About 1% of the contacts could not be determined as to whether they were symmetrical or asymmetrical. Of those 71.3% of the contacts that were asymmetrical, 36.5% of those were on dendrites **(Fig. 8D)** while 34.7% were on spines **(Fig. 8E)**. Of the 27.8% of the contacts that were symmetrical, 21.7% were on dendrites **(Fig. 8F)** and 6.1% were on spines.

Investigating synaptic contacts outside the implant area (located ∼0.25 mm lateral of the implant site), of the percentage of SC121+ labeled nerve terminals contacting SC121 negatively labeled postsynaptic dendrites (i.e., from the host), 90.6% were making an asymmetrical synaptic contact while 9.4% were making a symmetrical contact (total of 32 observations). Of the 90.6% making an asymmetrical synaptic contact, 79.3% were contacting spines while 20.7% were contacting dendrites. There were also nerve terminals from the host striatum (i.e., SC121 negative) contacting SC121+ dendrites. Of those contacts, 52.6% were asymmetrical and 47.4% were making a symmetrical contact (total of 19 observations) **(Suppl. Fig. 6A-D)**.

### hNSCs Improve Some Altered Intrinsic and Synaptic Membrane Properties of MSNs in Host zQ175 Mice

Given that we have previously shown that excitatory and inhibitory inputs to striatal MSNs and cortical pyramidal neurons in the zQ175 mouse model are altered (Indersmitten et al., 2015), we obtained whole-cell voltage clamp recordings to measure membrane and synaptic properties of neighboring host MSNs to determine whether transplanted hNSCs conferred modulatory outcomes **(Fig. 9A-D)**. Data from WT mice implanted with hNSCs and injected with vehicle only were pooled as there were no consistent differences in measures from MSNs between the two groups **(Table ID)**. We observed an improvement in the cell membrane properties of zQ175 MSNs from hNSC-implanted mice compared to zQ175 MSNs from mice receiving the vehicle only **(Fig. 9A)**. Previously, we showed MSNs from symptomatic zQ175 mice have higher membrane input resistances than MSNs from WT mice (Indersmitten et al., 2015). Compared with WT MSNs (hNSC-implanted and vehicle-injected, combined), zQ175 MSNs from vehicle-injected mice had significantly higher input resistances (WT 91.6±10.1 versus Q175-Veh 170.2±19.7 MΩ, p<0.001; Kruskal-Wallis ANOVA of ranks, followed by Holm-Sidak pairwise comparisons). Although input resistances were slightly higher in zQ175-hNSC MSNs (120.1±13.1 MΩ) compared to WT MSNs, this difference was not statistically significant demonstrating improvement in this electrophysiological property. Input resistances were significantly lower in MSNs from hNSC-implanted zQ175 mice compared to MSNs from vehicle-injected mice (p=0.04). There were no significant differences in cell membrane capacitances or membrane time constants across all three groups **(Fig 9A)**.

**Figure 9:**
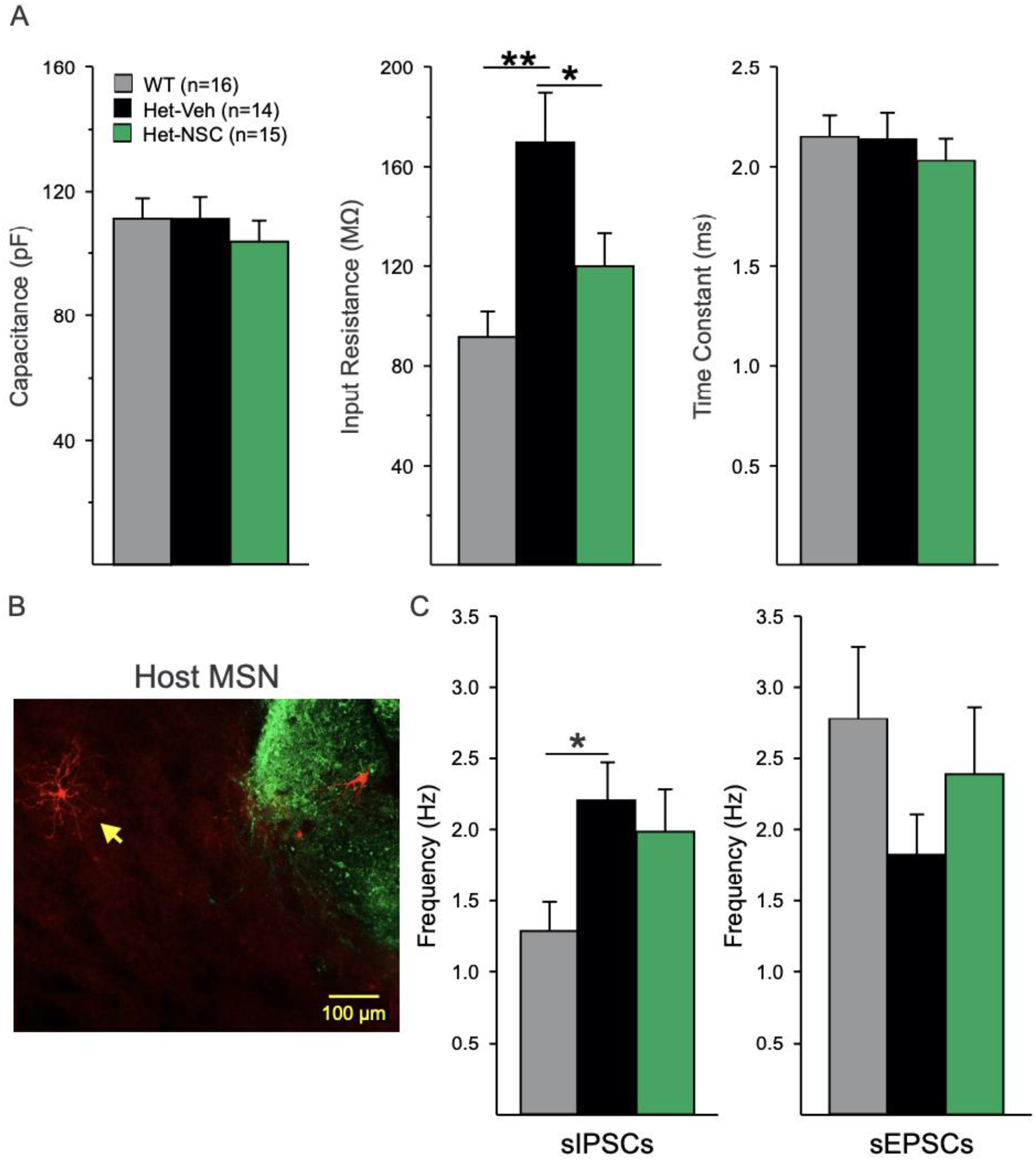
Rescue of some electrophysiological alterations. **A**. Cell membrane properties were recorded at a holding potential of −70 mV. **B**. Fluorescent image of a recorded and biocytin-filled MSN [yellow arrow pointing to filled neuron (red)] near SC121 immunostained cells and processes (green). **C**. Effects on inhibitory and excitatory synaptic activity. Statistical significance was measured between groups using Kruskal-Wallis one-way ANOVA on ranks followed by Holm-Sidak pairwise comparisons, In **A**. p<0.001 for WT versus Het-Veh (**) and p=0.04 for Het-Veh versus Het-NSC (*). In **C**. p=0.047 for WT versus Het-Veh (*).

In terms of hNSCs effects on synaptic activity, as reported previously, in zQ175 MSNs (Indersmitten et al., 2015) the frequency of sIPSCs recorded at a holding potential of +10 mV was increased compared to WT MSNs **(Fig. 9C)**. The increase was statistically significant in MSNs from Q175 mice injected with vehicle (WT 1.3±0.2 versus Q175-Veh 2.2±0.3 Hz, p=0.047; Kruskal-Wallis ANOVA of ranks, followed by Holm-Sidak pairwise comparisons) and although the sIPSC frequency was slightly higher in MSNs from hNSC-implanted zQ175 mice (2.0±0.3 Hz) compared to WT MSNs, it was not statistically significant (p=0.114) **(Fig. 9C)** demonstrating that the transplant reduced the increase in sIPSCs in MSNs from the zQ175 mice compared to WT mice. From the same cells, we recorded sEPSCs at a holding potential of −70 mV and in the presence of a GABA_A_ receptor antagonist (BIC, 10 µM). We observed a trend for decreased sEPSC frequency in MSNs from zQ175-Veh mice (1.8±0.3 Hz) compared to WT MSNs (2.8±0.5 Hz). Similarly, there was a trend for an increase in the frequency of sEPSCs in MSNs from the hNSC transplanted zQ175 mice (2.4±0.5 Hz).

Taken together, these data show long-term survival and differentiation of hNSCs into mainly neuronal lineages including a subset of mature-like MSNs and interneurons. The engrafted human cells establish connections with the host neurons and rescue specific electrophysiological and behavioral pathologies.

## Discussion

A major challenge in regenerative medicine approaches to treat neurodegenerative diseases, including HD, is enabling long-term assessment of cell fate, functional properties and potential rescue of disease-associated phenotypes (Jia et al., 2020; Kim et al., 2020). Here we evaluated whether hNSCs implanted in the striata of zQ175 mice are viable and integrate into the host tissue at a time point of 8 months after grafting (equating to roughly a third of the captive’s mouse 2-year lifespan). hNSCs survived and while most hNSCs had properties of immature neurons, approximately 20% of the cells evolved into more mature neurons with MSN- or interneuron-like properties and marker expression. Transplanted hNSCs receive synaptic inputs from neighboring cells or from the host, and innervate other hNSCs or host cells. Notably, grafted hNSCs increased striatal BDNF and pERK levels, reduced mHTT aggregated species, and ameliorated selected behavioral deficits of symptomatic zQ175 mice. Further, grafted cells modified host MSNs and rescued some of the altered membrane and synaptic properties observed in symptomatic mice. Thus, the mechanisms whereby implanted neurons rescue HD alterations appear to involve improved striatal MSN membrane properties and circuit connectivity, BDNF production, and prevention of the accumulation of mHTT aggregating species

To the best of our knowledge, this is the first study to examine, long-term (up to 8 months), electrophysiological and morphological properties of grafted hNSCs in a genetic mouse model of adult-onset HD. Also, it is the first to demonstrate that mature hNSCs evolve into the main types of resident neuronal populations in the striatum including not just MSNs, but also GABAergic and cholinergic interneurons. This is important as a comprehensive reconstruction of the striatal circuitry requires not just the presence of MSNs but a wide variety of interneurons which also are significantly affected in HD (Cepeda et al., 2013; Reiner et al., 2013; Holley et al., 2015; Holley et al., 2019a; Holley et al., 2019b).

Previously we demonstrated that hNSCs are viable and integrate into the host striatum in a severe model of HD, the R6/2 (Holley et al., 2018; Reidling et al., 2018). However, due to rapid progression of the phenotype in these mice, examination of transplanted hNSCs was limited by time, about 4-6 weeks after implantation. We also reported data from the long-lived homozygous Q140 HD mouse model (Reidling et al., 2018), however, cell survival was not optimal. This most likely occurred due to insufficient immunosuppression methods, and measurements of cell differentiation or electrophysiology and connectivity visualized by EM were not feasible. Here we show that hNSCs can survive 8 months after injection, integrate into the host striatum, and improve some of the abnormal MSN membrane and synaptic properties of the heterozygous zQ175 HD model. Further, a subset of hNSCs become more mature and display properties similar to those of MSNs and some types of interneurons. EM and electrophysiology also suggest connections from the host may originate in the cortex and support a potential reconnection of the corticostriatal pathway that is lost during HD progression and that reconnection could contribute to restoration of normal motor and cognitive functions. We hypothesize that the higher frequencies of synaptic inputs are due to the maturation and integration of hNSCs within the host tissue, as we did not observe hNSCs with these synaptic properties in our previous study using R6/2 hNSC-implanted mice. Unlike the R6/2 mice, MSNs from zQ175 mice do not exhibit epileptiform activity in the presence of BIC. The R6/2 mouse model demonstrates cortical hyperexcitability that can be exacerbated pharmacologically when inhibition is reduced (Cummings et al., 2009). This is reflected in R6/2 MSNs as large amplitude excitatory events followed by a barrage of high frequency, small amplitude events. We did not observe this electrophysiological phenotype in any of the zQ175 or WT MSNs. Electrophysiologically, ∼80% of hNSCs remained “immature”, despite surviving in the graft for 8 months. It is unknown why these cells do not differentiate. It is possible that with longer implantation time these cells also become more mature. We previously showed that transplanted hNSCs in R6/2 mice exhibited evidence of neuron-restricted progenitor markers DCX, Beta-III tubulin, and MAP-2 (Reidling et al., 2018). As hNSCs typically take several months to terminally differentiate during development, we expected to observe further differentiation of transplanted cells in the zQ175 8-month implant cohort. Interestingly, although we found this to be the case with some cell staining for the post-mitotic neuronal marker NeuN, as well as interneuron and MSN markers, we still observed clusters of ESI-017 hNSCs to be DCX positive after 8 months. Perhaps the DCX positive hNSCs have not received the signals to differentiate as many seem to be on the interior of the implant. It is important to note that the hNSCs appear to have received signals to stop proliferating (no Ki67 or EdU incorporation) and start down a path to differentiate (loss of nestin). Even though most cells did not fully differentiate and did not become more mature, they still looked healthy and displayed neuronal properties, including the capacity to generate action potentials.

Although we can conclude that about 20% of the large hNSCs display physiological features of mature neurons, it is difficult to determine the specific cell-types based solely on their electrophysiological properties. Supporting evidence was provided by IHC and the presence of specific striatal neuronal markers. MSN-like hNSCs displayed Ca^2+^ currents typically observed in mature MSNs and IHC demonstrated the presence of DARPP-32 and Ctip2, which label striatal MSNs. Other hNSCs fired rhythmically, had increased input resistances (compared to MSNs), lacked characteristic Ca^2+^ currents, and received rhythmic GABAergic synaptic events, suggesting that these cells could be GABAergic interneurons. In support, IHC from these grafts indicated the presence of GAD and CR, specific interneuron markers. Some large hNSCs also displayed rhythmic bursting and low-threshold spikes, reminiscent of the somatostatin-expressing (LTS) interneurons (Tepper et al., 2018; Holley et al., 2019b). This observation is significant in terms of HD because these two interneuron subtypes are spared during disease progression. In fact, CR+ interneurons appear increased in HD (Cicchetti and Parent, 1996), suggesting that they could have neuroprotective properties or alternatively they are selected for within the HD niche. Indeed, both somatostatin and calretinin have been shown to be neuroprotective (Kumar, 2008) and a study on grafted fetal striatal tissue demonstrated the presence of graft-derived neurons expressing DARPP-32, calretinin and somatostatin (Capetian et al., 2009). Another possibility is that some large hNSCs could have evolved into NPY interneurons. Interestingly, in HD, MSNs expressing NPY are spared and their numbers are even upregulated in HD patients (Wagner et al., 2016). Based on passive and active membrane properties, some cells also differentiated into large cholinergic interneurons. In contrast, it seems unlikely that differentiated hNSCs became fast-spiking interneurons, at least in a mature state.

Results of cell fate in our HD mouse studies uniquely appear to follow a neuronal developmental path in contrast to a more gliogenic outcome, potentially due to the differentiation potential of the starting material or the transplantation niche (Goldberg et al., 2017; Qian et al., 2020; Yoon et al., 2020). A recent study using a rat model of HD induced by intrastriatal quinolinic acid injection, showed that human embryonic stem cell-derived MSN progenitors differentiate *in vitro*, undergo maturation, integrate into host circuits, and display properties similar to those of the host striatum 2 months after transplantation (Besusso et al., 2020), suggesting feasibility of transplanting differentiated MSN progenitors. Notably, behavioral studies in this model demonstrated functional recovery of some impaired sensorimotor responses but not in more complex behaviors (e.g., rotarod test). Some cells were proliferative, the yield of DARPP32/Ctip2 double-labeled MSNs was relatively low, and there was some degree of contamination from cortical neurons. However, it is not known whether excitotoxicity models reflect the biochemical environment of the genetic mutation. Interestingly in that study, similar to ours, a low percentage of grafted cells expressed interneuronal markers (calbindin and calretinin), however these cells were not characterized electrophysiologically. Other strategies to generate striatal neurons are under investigation. For example, a recent study used an *in vivo* cell conversion technology to reprogram striatal astrocytes into GABAergic neurons through AAV-mediated ectopic expression of NeuroD1 and Dlx2 transcription factors (Wu et al., 2020). The striatal astrocyte-converted neurons showed action potentials and synaptic events, and projected their axons to the appropriate target regions. Behavioral analyses of these treated R6/2 mice showed a significant extension of life span and improvement of motor deficits (Wu et al., 2020).

In conclusion, our studies support future development of stem cell-based therapies. While mHTT/HTT-lowering strategies and gene editing are promising as therapies and are in various stages of clinical trials, a pressing issue is how to replace the striatal cell loss occurring in HD patients even prior to overt symptomatic onset. The present results support our previous findings that implanted cells may provide nursing effects through enhanced BDNF levels and reduction of pathological mHTT. Importantly, hNSCs establish synaptic contacts with host cells and among themselves, differentiate into a wide variety of striatal resident cells, and form the building blocks for circuit regeneration. Given the electrophysiological and EM results presented here, there is also promise that over time, transplanted cells can make beneficial synaptic connections and replace lost functions. Thus, our preclinical study demonstrates that hNSCs transplanted into a relevant HD model brain survive for long periods of time and may potentially be utilized for restoration of circuity and cell replacement in the clinic.

## Methods

### Mice

All experimental procedures were in accordance with the Guide for the Care and Use of Laboratory Animals of the NIH and animal protocols were approved by Institutional Animal Care and Use Committees at the University of California Irvine (UCI), the University of California Los Angeles (UCLA), and the Portland VA Medical Center, AAALAC accredited institutions. zQ175 heterozygous (Het) mice and their wildtype (WT) littermates were obtained from breeding colonies maintained at UCI (zQ175 Het mice had ∼163-199 CAG repeats, Laragen, Culver City, CA). All mice were housed on a 12/12-hr light/dark schedule with *ad libitum* access to food and water. Mice were group-housed as mixed treatment groups and only males were single-housed for the running wheel test. Groups included: 10 male zQ175 Het hNSC, 8 female Het hNSC, 9 male zQ175 Het vehicle, 9 female zQ175 Het vehicle, 7 male WT hNSC, 7 female WT hNSC, 6 male WT vehicle, 6 female WT vehicle.

### Cells

The use of hESCs and hNSCs was approved by Human Stem Cell Research Oversight Committees (hSCRO) at UCI, UCLA, and UC Davis. ESI-017 is one of the six clinical-grade hESC lines generated from supernumerary embryos by the Singapore Stem Cell Consortium (Crook et al., 2007). Their use for therapeutic application adheres to US FDA regulations for use of human cells. Of those lines, four (including ESI-017) were chosen for the generation of Good Medical Practice (GMP) hESC banks for preclinical research based on the absence of human and non-human pathogens (Crook et al., 2007; Sivarajah et al., 2010). Subsequently, an hNSC line was differentiated from the GMP-grade hESC line ESI-017 as described previously (Reidling et al., 2018). ESI-017 hNSCs were acquired as frozen aliquots, thawed, and then cultured for a minimal time out of thaw (2-3 days) using the same media reagents as the GMP facility prior to dose administration. The cells were not passaged.

### Surgery

Two and a half month-old zQ175 Het mice and WT littermates were anesthetized, placed in a stereotaxic frame and injected with either 100,000 hNSCs per side (2 μl/injection) or vehicle (2 μl HBSS with 20 ng/ml epidermal growth factor [STEMCELL Technologies, #78003] and human fibroblast growth factor [STEMCELL, #78006]) as a control treatment using a 5 μl Hamilton microsyringe (33-gauge) and an injection rate of 0.5 μl/min. Coordinates relative to Bregma were AP: 0.00, ML: +/- 2.00, and DV −3.25 mm. For immunosuppression, all mice received IP injections of cyclosporine (10 mg/kg, daily thereafter) and mouse CD4 Ab (10 mg/kg, weekly thereafter) the day before surgery and continued until mice were sacrificed (8 months after implantation).

### Biochemical, Molecular, and Immunohistochemical Analyses

Four male mice per group were given IP injections of EdU (Thermofisher Scientific) 24 hr prior to sacrifice. Mice were euthanized by pentobarbital overdose and perfused with 0.01 M PBS. Striatum and cerebral cortex were dissected out of the left hemisphere and flash-frozen for biochemical analysis. The other halves were post-fixed in 4% paraformaldehyde, cryoprotected in 30% sucrose, and cut at 40 μm on a sliding vibratome for immunohistochemistry (IHC). Sections were rinsed three times and placed in blocking buffer for 1 hr (PBS, 0.02% Triton X-100, 5% goat serum), and primary antibodies placed in block overnight (ON) at 4°C. Sections were rinsed, incubated for 1 hr in Alexa Fluor secondary antibodies, and mounted using Fluoromount G (Southern Biotechnology). Primary antibodies used include: Anti-Ki67 (Abcam, ab16667), Anti-Ku80 (Abcam, ab80592), Anti-Nestin (Millipore Sigma, MAB5326), Anti-GFAP (Abcam, ab4674), Anti-DCX (Fisher Millipore, AB2253MI), Anti-NeuN (Abcam, ab177487), Anti-Calretinin (Abcam, ab16694), Anti-vGlut1 (Abcam, ab180188), Anti-HNA (Abcam, ab191181), Anti-GAD65/67 (Abcam, ab49832), Anti-BetaIII tubulin (Abcam, ab78078), Anti-DARPP32 (Abcam, ab40802), Anti-Ctip2 (Abcam, ab233713). For IHC a minimum of four mice per group were analyzed. *DAB staining for ChAT:* Sections were rinsed three times then 30 min in 3% H_2_O_2_ and 10% Methanol rinsed and placed in blocking buffer for 1 hr (TBS + 5% normal rabbit serum (NBS Vector S-5000) + 0.1% TritonX-100), then primary antibody (goat anti-ChAT 9 Millipore AB144P) placed in block overnight at 4°C. Sections were rinsed, incubated for 1 hr in secondary antibody (rabbit anti-goat biotinylated secondary), incubated in ABC solution (Vector PK-6100) for 1 hr at RT then 1-3min in DAB, rinse and mount tissue on slide. *Confocal Microscopy:* Sections were imaged with Bio-Rad Radiance 2100 confocal system using lambda-strobing mode. Images represent either single confocal z-slices or z-stacks. *Whole cell tissue lysis:* Lysis was performed in RIPA buffer supplemented with protease inhibitors (Complete Mini, Roche Applied Science), 0.1 mM PMSF, 25 mM NEM, 1.5 mM aprotinin, and 23.4 mM leupeptin by douncing then sonicated for 10 seconds, 3 times at 40% amplitude on ice. Samples were quantified using Lowry protein assay. *Soluble/Insoluble Fractionation:* Striatal tissue was processed as described previously (Ochaba et al., 2016). *Western analysis:* RIPA lysates were resolved by reducing and running 60µg of protein on 4-12% Bis-Tris Poly-Acrylamide gels (PAGE). Antibodies: Anti-BDNF (Santa Cruz Biotechnology, clone N-20, for mature BDNF, cat.no.sc-546), Anti-ERK (Cell Signaling Technology, cat.no. 9102), Anti-pERK (Cell Signaling Technology, cat.no. 9106), Anti-alpha tubulin (Sigma-Aldrich, cat.no. T6074). Quantification of bands was performed using software from the NIH program ImageJ and densitometry application. 50µg of reduced, insoluble protein from Insoluble Fractions were resolved on 3-8% Tris-Acetate Poly-Acrylamide gels. Membranes were blocked in Starting block (Invitrogen) for 20 minutes at room temperature and probed in primary antibody overnight at 4°C. Insoluble protein was quantified as relative protein abundance as previous (Ochaba et al., 2016). Antibodies: Anti-HTT (Millipore, #MAB5492; RRID: AB_347723).

### Behavioral Tests

Males and females were used except for the running wheel, where only males were used since estrus cycle influences running activity. Genotypes or treatments were unknown to the experimenter. All tests were done during the light phase except for the running wheel, where mice were allowed 24 hr free access to the task. Running wheel data are only described for the dark phase. Slope of motor learning was calculated as mean nightly running wheel rotations per 3 minutes on night 5 minus night 2 divided by total number of nights (3) for initial and night 13 minus night 2 divided by total number of nights (11) for overall. Behavioral tasks, running wheel and open field, were performed in a manner to those previously described (Hickey et al., 2008; Reidling et al., 2018).

### Electrophysiology

For electrophysiological studies we used 12 female mice (10.5 month-old) shipped live to UCLA from UCI. Groups included: 4 zQ175 Het hNSC, 4 zQ175 Het vehicle, 2 WT hNSC, 2 WT vehicle. Mice were anesthetized and transcardially perfused with high sucrose-based slicing solution. Coronal slices (300 μm) were transferred to an incubating chamber containing standard artificial cerebrospinal fluid (ACSF). MSNs and hNSCs were visualized using infrared illumination with differential interference contrast optics (IR-DIC). All recordings were performed in or around the injection site (recorded MSNs were adjacent to the graft, ∼150-250 µm). Biocytin (0.2%) was added to the patch pipette for cell visualization and location of recorded cells. Spontaneous postsynaptic currents were recorded in the whole-cell patch clamp configuration in ACSF. Membrane currents were recorded in gap-free mode. Cells were voltage-clamped at +10mV and spontaneous inhibitory postsynaptic currents (sIPSCs) were recorded in ACSF. Spontaneous excitatory postsynaptic currents (sEPSCs) from grafted cells were recorded in ACSF at −70 mV (baseline). sEPSCs from MSNs, were recorded in the presence of the GABA_A_ receptor blocker, bicuculline methobromide (BIC, 10 µM, Tocris, Minneapolis, MN) to better isolate glutamatergic events. Spontaneous synaptic currents were analyzed using the MiniAnalysis software (version 6.0, Synaptosoft, Fort Lee, NJ). To evoke responses in grafted cells, we used a monopolar glass electrode (impedance 1 MΩ) which was placed 200-300 μm lateral to the graft. Following recordings, slices were fixed with 4% PFA, then transferred to 30% sucrose at 4°C until IHC processing. To identify biocytin-filled recorded cells and hNSCs, fixed slices were washed, permeabilized with triton (0.7%) and blocked for 4 h, followed by incubation with SC121 (1:1000). After washing, slices were incubated in goat, anti-mouse Alexa-488 (1:1000, Life Technologies, Carlsbad, CA Catalog #:A-11001) and streptavidin conjugated with Alexa-594 (1:1000, Life Technologies Catalog #: S11227). Slices were washed, mounted, and cells visualized with a Zeiss Apotome confocal microscope.

### Electron Microscopy (EM)

Female zQ175 mice implanted with hNSCs for 8 months (n=3 per group) at UCI were sent live to the Portland VA Medical Center. Mice were anesthetized and perfused with EM fixative (2.5% glutaraldehyde, 0.5% paraformaldehyde, and 0.1% picric acid in 0.1 M phosphate buffer [pH 7.4]). Brains were then collected and further processed in a Pelco Biowave Pro+ (Ted Pella, Inc, Redding, CA), as previously reported (Moore et al., 2020), and then washed in PBS and stored overnight at 4° C. Striatum containing hNSCs (equivalent to +1.4 to +0.14 mm from Bregma) (Franklin and Paxinos, 2007) was cut at 60 μm using a vibratome (Leica Microsystems). After pre-embed IHC of the striatum using diaminobenzidine (DAB) (Sigma, St Louis, MO) or ImmPACT® VIP Substrate, Peroxidase (HRP) (Cat #: SK-4605)(Vector Labs, Burlingame, CA), hNSC antibody (SC121, 1:100; Takara: Cat #: Y40410) and vGlut 1 (vesicular glutamate transporter 1) antibody (1:000; Synaptic Systems, Germany, Cat #: 135-303), the tissue was processed for EM as previously described (Walker et al., 2012; Parievsky et al., 2017; Reidling et al., 2018; Moore et al., 2020). Two striatal slices were selectively double labeled for SC121 (DAB) and vGlut1(VIP) to determine if vGlut1 labeled terminals originating from the cortex (Kaneko et al., 2002) innervated the implantation site. Striatal slices were embedded flat between two sheets of ACLAR (Electron Microscopy Sciences, Hatfield, PA) overnight in a 60°C oven to polymerize the resin. The area containing hNSCs was micro-dissected from the embedded slice and superglued onto a block for thin sectioning. Photographs were taken on a JEOL 1400 transmission electron microscope (JEOL, Peabody, MA) of DAB labeled structures (i.e., SC121) and for a small number of sections, double labeled with DAB (SC121) and VIP (vGlut1)-labeled structures (i.e., hNSC-labeled cells, dendrites, nerve terminals). The DAB labeled structures (i.e., SC121) were photographed both within and located ∼0.25 mm outside of the implantation zone, at a final magnification of 46,200 using a digital camera (AMT, Danvers, MA). Since the DAB and DAB/VIP labeling was restricted to the leading edge of the thin-sectioned tissue, only the area showing DAB and DAB/VIP labeling was photographed. The percentage of SC121/DAB-labeled asymmetrical and symmetrical synaptic contacts onto dendrites and spines within and outside the implant area was quantified.

### Statistical Analysis

Results are from a single cohort except for Western blots for BDNF and pERK/ERK and IHC for ChAT, which were from a different subset. Numbers were determined to have sufficient power using an analysis prior to the study. Assessment of differences in outcome were based upon previous experience and published results (Hockly et al., 2003; Hickey et al., 2005) for HD models, and applying power analysis (G Power [http://www.psycho.uni-duesseldorf.de/abteilungen/aap/gpower3/]) led us to a minimal n=5 for behavior and n=3 for biochemical analysis. Statistical significance was achieved as described using rigorous analysis. All findings are reproducible. Experiments were performed at least 3 times using at least 3 different mice (biological replicates) and in specific cases tissue from one mouse was used multiple times (technical replicates); for example, in IHC at least 3 different mouse brains were used and multiple sections from each brain were examined to obtain data. Multiple statistical methods are further detailed above, in figure legends, or in Supplementary Experimental Procedures. Since the EM data are based on 3 implanted zQ175 mice, and not comparing them against the WT mice, the percentages reported are a comparison within a single group of 3 experimental mice, therefore, there was no statistical comparison. Statistical tests for behavioral tasks used one-way ANOVA followed by Tukey’s HSD test with Scheffé, Bonferroni, and Holm multiple comparison *post hoc* methods. Data met the assumptions of the statistical tests used, and p values <0.05 were considered significant. All mice were randomly assigned and tasks performed in a random manner with individuals blinded to genotypes and treatment. Statistical comparisons of densitometry results were performed by one-way ANOVA followed by Tukey HSD and Bonferroni *post hoc* tests. For electrophysiology data, all statistical analyses were performed using SigmaPlot 13.0 software. Differences between multiple group means were assessed with appropriate one-way ANOVAs followed by Bonferroni *post hoc* tests, Kruskal-Wallis one-way ANOVA on ranks followed by Holm-Sidak *post hoc* tests or Student’s *t*-tests (unpaired) when only two groups were compared. Significance levels in the figures are given as specific p-values and data are expressed as mean ± SEM.

## Acknowledgments

Funding was provided by the California Institute for Regenerative Medicine (CIRM ETAII TR2-01841), and NIH grants NS096994 and U54HD087101 (MSL).

We thank BioTime, Inc and AgeX for the ESI-017 cell line and the UC Davis Flow Cytometry Shared Resource, 2921 Stockton Blvd., Suite 1300/1670 Sacramento, CA 95817 for flow analysis. We also thank the UCI Institute for Memory Impairments and Neurological Disorders, the Sue and Bill Gross Stem Cell Center and the Optical Biology Shared Resource of the Cancer Center Support Grant (CA-62203) at the University of California, Irvine for facilities and assistance in carrying out studies.

## Author contributions

J.C.R., S.M.H., C.C., M.S.L., L.M.T. designed experiments and analyzed data. J.C.R., S.M.H., C.C., L.M.T., M.S.L. wrote the manuscript. S.Y., A.L., I.O. performed experiments in mice, A.L. and M.N. did IHC and E.S.M. and J.C.R. performed analyses. S.M.H., C.C. performed electrophysiology and analyzed data with guidance from M.S.L. C.M. performed EM with guidance from C.K.M. L.K. cultured hNSCs at UCI. B.F., D.C.-B. and G.B. supplied ESI-017 hNSCs and characterizations from GMP facility at UCD.

## Supplementary Figures

**Supp Figure 1:**
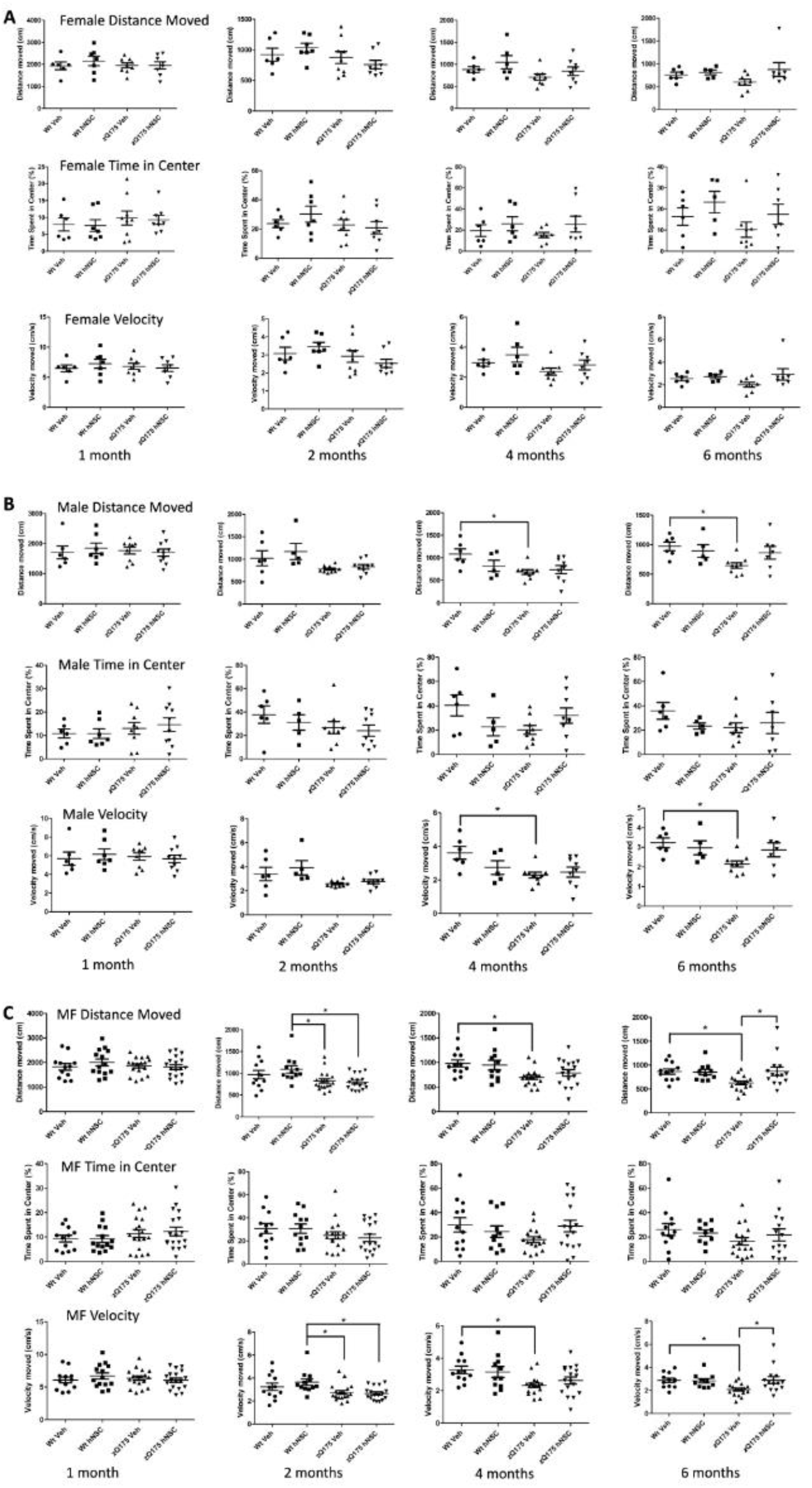
Open Field behavior. Total distance traveled in the open field 6 months post implant. Mice were subjected to the open field and total distance in centimeters of their respective tracks were combined and statistically analyzed to visualize any differences in ambulation. Time spent in center was also measured. In addition, velocity traveled in centimeters/sec of their respective tracks were combined and statistically analyzed to visualize any differences in time of ambulation. **A**. Females **B**. Males and **C**. Males and Females combined. Groups for open field at 1 month included: 10 male zQ175 Het hNSC, 8 female Het hNSC, 9 male zQ175 Het veh, 9 female zQ175 Het veh, 7 male WT hNSC, 7 female WT hNSC, 6 male WT veh, 6 female WT veh. Groups for open field at 6 months included: 7 male zQ175 Het hNSC, 7 female Het hNSC, 9 male zQ175 Het veh, 8 female zQ175 Het veh, 5 male WT hNSC, 5 female WT hNSC, 6 male WT veh, 6 female WT veh. Results are expressed as the mean ± S.E.M with one-way ANOVA Bonferroni post test: *In order of graphs p=0.03, p=0.04, p=0.03, p=0.04, p=0.01, p=0.01, p=0.01, p=0.01, p=0.01, p=0.01.

**Suppl Figure 2:**
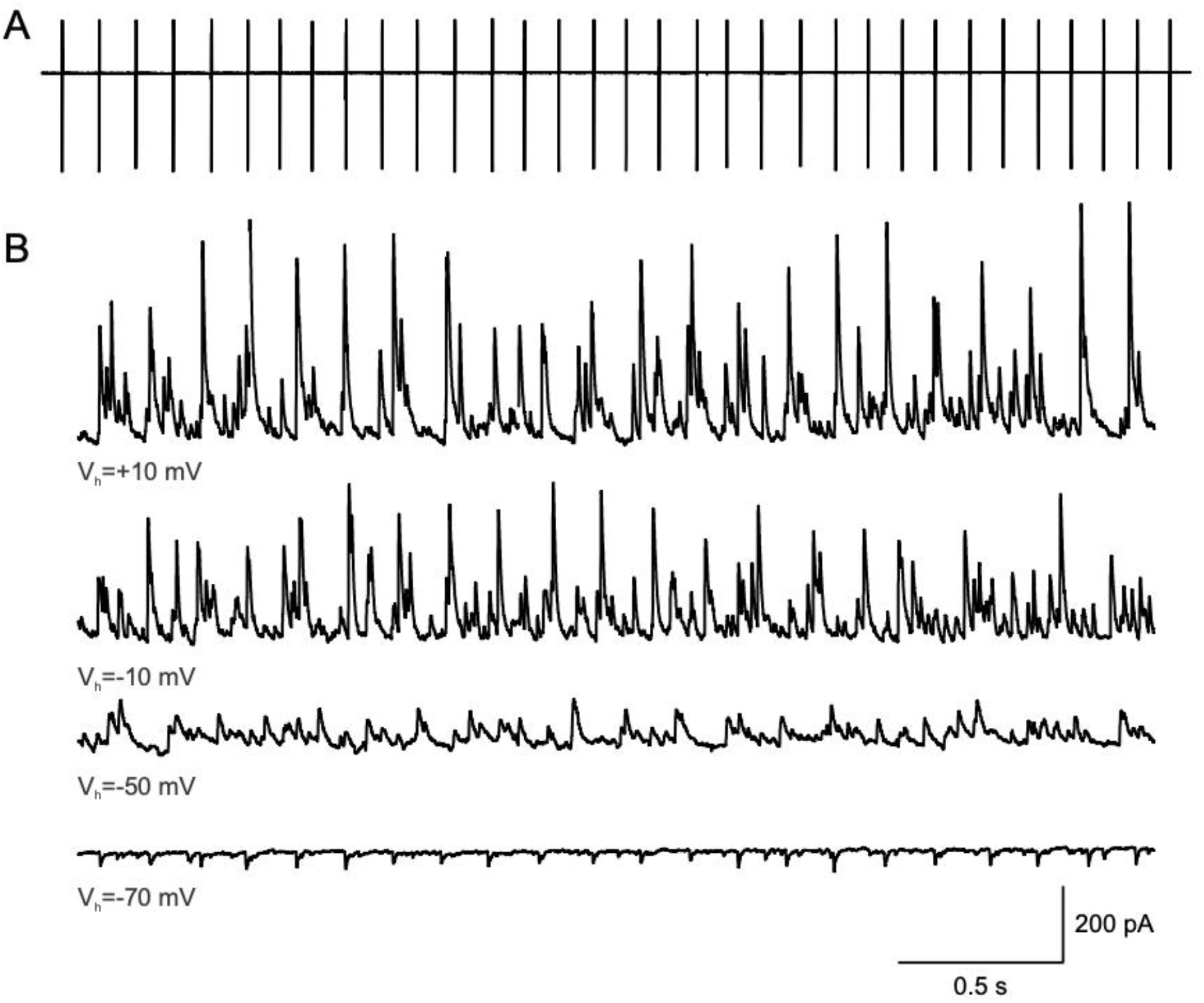
hNSCs display rhythmic activities. **A**. In cell-attached mode this cell from a zQ175 mouse displayed autonomous, rhythmic firing activity. **B**. In voltage clamp mode, spontaneous, rhythmic synaptic events can be observed at different holding potentials (bottom 4 traces). Spontaneous synaptic events were tentatively assumed to be GABAergic since the reversal potential for GABA occurred at ∼-60 mV. This type of activity is not observed in normal conditions in striatal MSNs and this cell was assumed to be interneuron-like.

**Suppl Figure 3:**
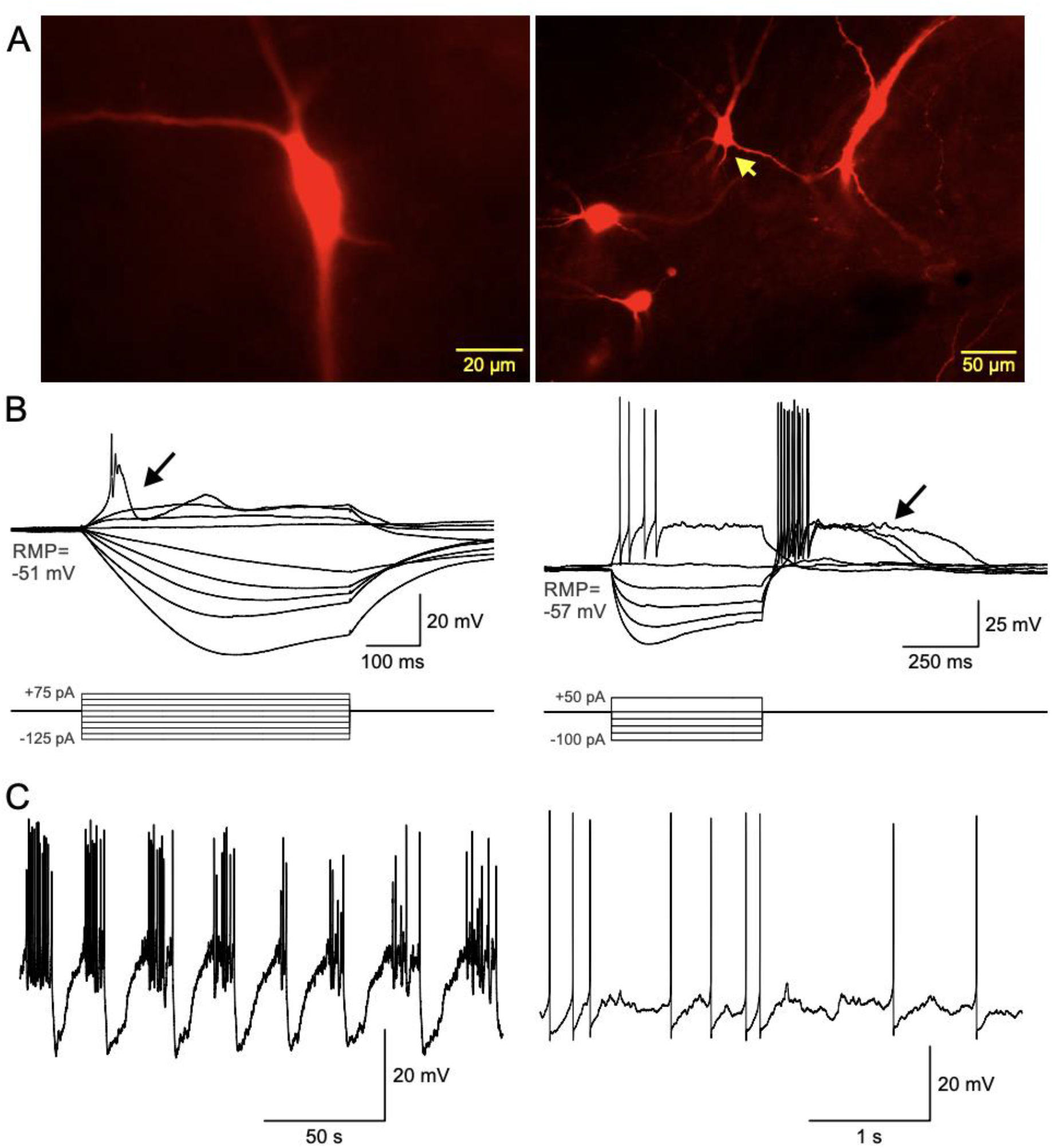
A. Large hNSCs from two different zQ175 implanted mice were recorded and filled with biocytin. The hNSC on the left panel and the cell with the yellow arrow on the right panel displayed LTS-like interneuron properties. **B**. Traces are electrophysiological recordings in current clamp mode. Both cells displayed prominent inward rectification and low-threshold Ca^2+^ spikes (arrows). **C**. Upon hyperpolarization by negative current injection the cell on the left displayed rhythmic membrane oscillations and bursts of action potentials. The interneuron-like hNSC on the right displayed low-threshold Ca^2+^ spikes and fired spontaneously at resting membrane potential (−57 mV). Both cells shared similarities with striatal LTS (somatostatin-expressing) interneurons.

**Suppl Figure 4:**
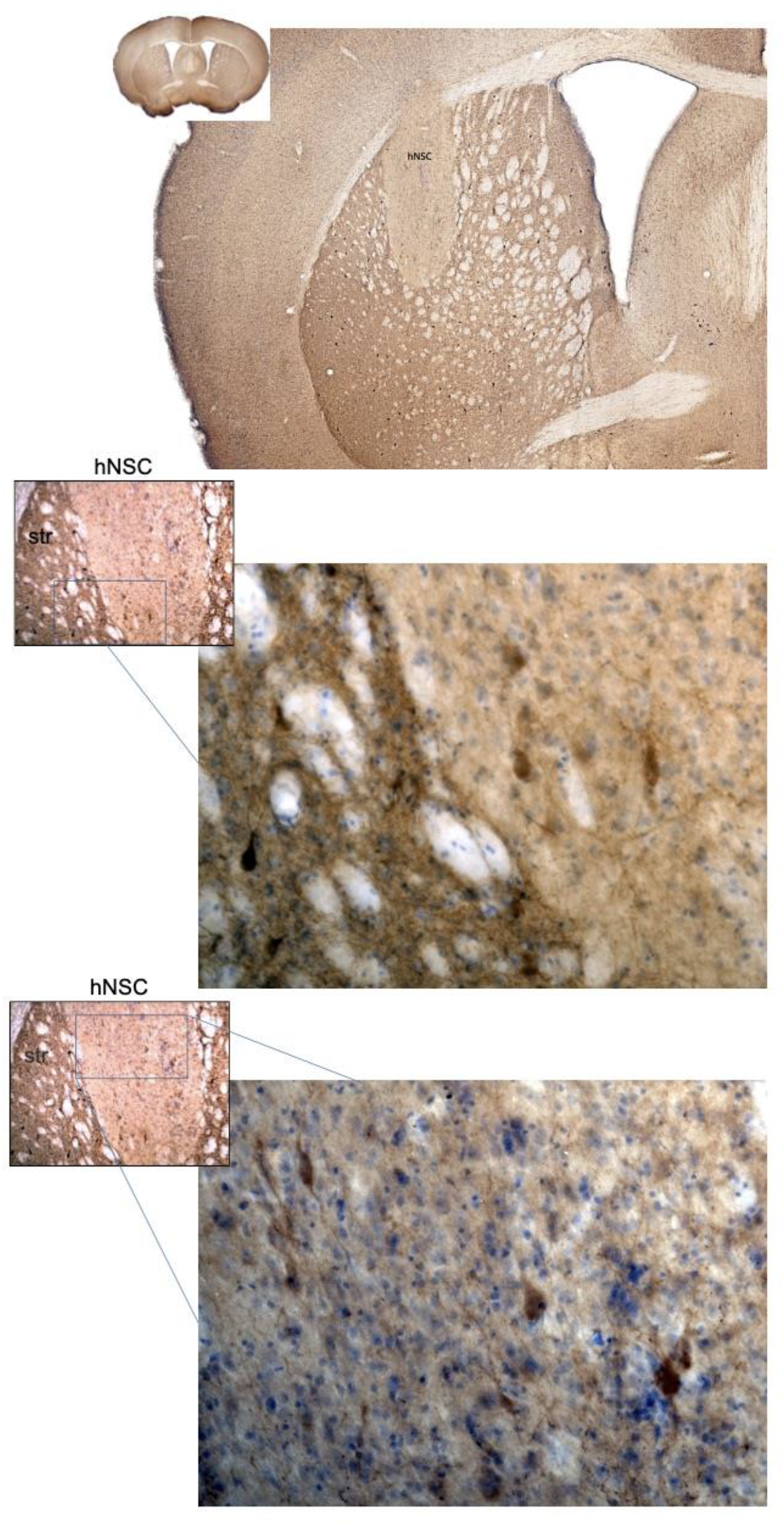
IHC for ChAT demonstrated that some hNSCs express the cholinergic marker. 5x mag. of hNSCs in zQ175 showing overall implant site. 10X mag. then box indicating 20x mag. showing area in and around implant site that has DAB positive (brown) ChAT expressing cells.

**Suppl Figure 5:**
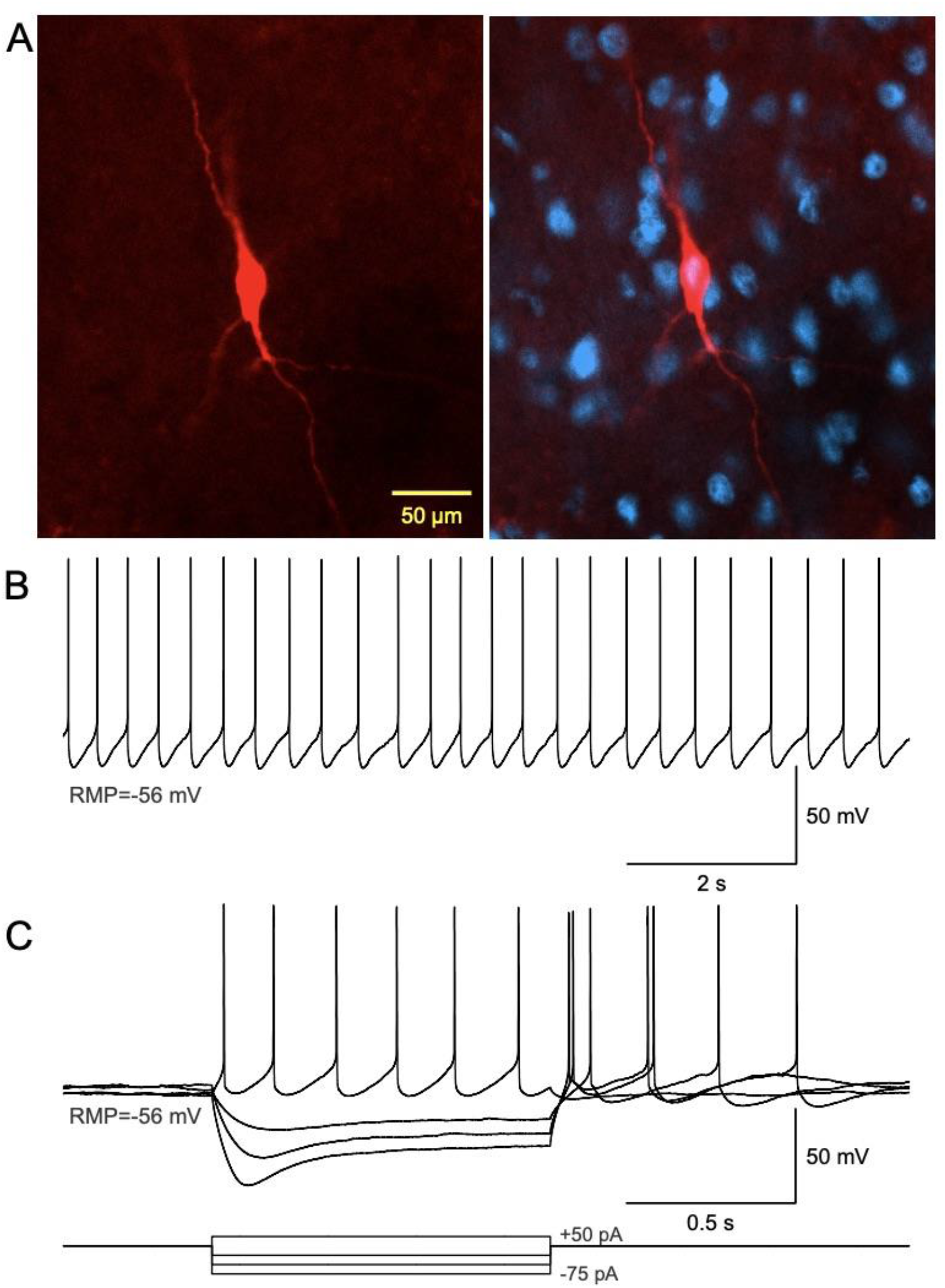
**A**. Left panel shows a biocytin-filled, large hNSC. The right panel illustrates co-localization of the human marker Ku80 (blue) and biocytin (red). **B**. In current clamp mode, this cell fired spontaneous, rhythmic action potentials (2-3 Hz), typical of striatal cholinergic interneurons. **C**. Hyperpolarizing the cell also demonstrated delayed inward rectification, another signature of striatal cholinergic interneurons.

**Suppl Figure 6:**
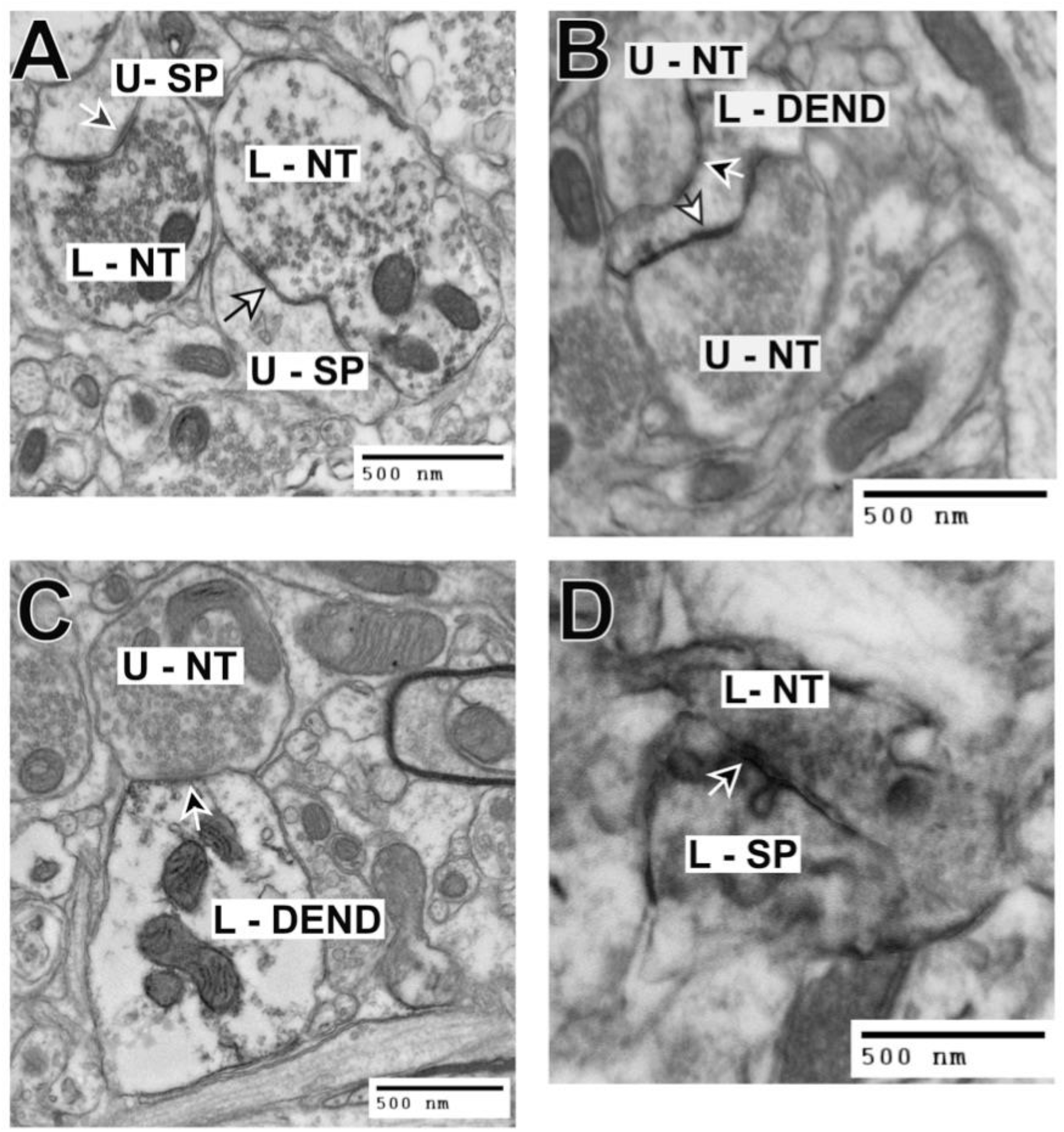
Striatal synaptic contacts located outside the stem cell implantation zone. **A**. A labeled nerve terminal (L-NT) is making either an asymmetrical synaptic contact (black arrow with white outline) or a symmetrical contact (white arrow with black outline) with an underlying hNSC negative/unlabeled dendritic spine (U-SP)(note the darkened DAB reaction product within the nerve terminals). The hNSC labeled nerve terminals originate from the implanted stem cells, while the unlabeled dendritic spines originate from the host striatum. **B**. Unlabeled nerve terminals (U-NT) are making either an asymmetrical synaptic contact (black arrow with white outline) or a symmetrical contact (white arrow with black outline) with an underlying hNSC positive/labeled dendrite (L-DEN) (note the darkened DAB reaction product within the dendrite). The hNSC labeled dendrite originates from the implanted stem cells, while the unlabeled nerve terminals originate from the host striatum. **C**. Unlabeled nerve terminal (U-NT) making an asymmetrical synaptic contact (arrow) with an underlying hNSC positive/labeled dendrite (L-DEND)(note the darkened DAB reaction product within the dendrite). The hNSC labeled dendrite originates from the implanted stem cells, while the unlabeled nerve terminal originates from the host striatum. **D**. Labeled nerve terminal (L-NT) is making a symmetrical synaptic contact (arrow) with an underlying hNSC positive/labeled dendritic spine (L-SP) (note the darkened DAB reaction product within the nerve terminal and spine). The hNSC labeled nerve terminal and dendritic spine originate from the implanted stem cells.

